# Hypercapnia limits β-catenin-mediated alveolar type 2 cell progenitor function by altering Wnt production from adjacent fibroblasts

**DOI:** 10.1101/2022.01.12.475264

**Authors:** Laura A. Dada, Lynn C. Welch, Natalia D. Magnani, Ziyou Ren, Patricia L. Brazee, Diego Celli, Annette S. Flozak, Anthea Weng, Mariana Maciel-Herrerias, Vitalii Kryvenko, István Vadász, Constance E Runyan, Hiam Abdala-Valencia, Masahiko Shigemura, S. Marina Casalino-Matsuda, Alexander V. Misharin, G.R. Scott Budinger, Cara J. Gottardi, Jacob I. Sznajder

**Author notes:** Corresponding author. Laura A. Dada, 303 E Superior St, Simpson-Querrey 5-405, Chicago, IL 60611, USA.

## Abstract

Persistent symptoms and radiographic abnormalities suggestive of failed lung repair are among the most common symptoms in patients with COVID-19 after hospital discharge. In mechanically ventilated patients with ARDS secondary to SARS-CoV-2 pneumonia, low tidal volumes to reduce ventilator-induced lung injury necessarily elevate blood CO_2_ levels, often leading to hypercapnia. The role of hypercapnia on lung repair after injury is not completely understood. Here, we show that hypercapnia limits β-catenin signaling in alveolar type 2 (AT2) cells, leading to reduced proliferative capacity. Hypercapnia alters expression of major Wnts in PDGFRa+-fibroblasts from those maintaining AT2 progenitor activity and towards those that antagonize β-catenin signaling and limit progenitor function. Activation of β-catenin signaling in AT2 cells, rescues the inhibition AT2 proliferation induced by hypercapnia. Inhibition of AT2 proliferation in hypercapnic patients may contribute to impaired lung repair after injury, preventing sealing of the epithelial barrier, increasing lung flooding, ventilator dependency and mortality.

## INTRODUCTION

As of this writing, more than 297 million people have been diagnosed globally with Coronavirus Disease 2019 (COVID-19) and nearly 5.5 million people have died (https://coronavirus.jhu.edu). Severe COVID-19 presents as acute respiratory distress syndrome (ARDS) where injury of the alveolar epithelial barrier causes flooding of the alveolar space and pulmonary edema which in severe cases may require mechanical ventilation(*1, 2*). In the US alone, there are currently 57 million survivors of COVID-19. Studies focused on post-acute sequelae of COVID-19 (PASC) suggest persistent respiratory symptoms while radiographic abnormalities and requirement for supplemental oxygen are common in survivors of COVID-19, particularly those requiring high flow oxygen therapy or mechanical ventilation (*3*). These persistent symptoms suggest that failure of normal lung repair mechanisms can prevent complete recovery of lung function in a substantial fraction of COVID-19 survivors, with significant public health impact.

Low tidal volume ventilation is a proven strategy to reduce the incidence and severity of ventilator induced lung injury (*4–6*). The alveolar hypoventilation associated with low tidal volumes necessarily induces increases in blood CO_2_ levels, described clinically as hypercapnia. Hypercapnia is exacerbated by increases in steady state carbon dioxide production and dead space ventilation common in patients with severe SARS-CoV-2 pneumonia and ARDS (*4, 6, 7*). We and others found that exposure to hypercapnia activates signaling pathways detrimental to alveolar epithelial wound healing and migration (*4–10*), but mechanisms by which hypercapnia impairs epithelial repair remain incompletely understood.

While baseline turnover in the alveolar epithelium is slow, alveolar injury results in rapid and robust alveolar type 2 (AT2) cell proliferation and differentiation to restore barrier function and gas exchange (*11–13*). During homeostasis and after lung injury, lineage tracing studies suggest that AT2 cells self-renew and serve as progenitor cells for alveolar type 1 (AT1) cells (*13–16*). Our group was among the first to report that Wnt-β-Catenin (βcat) signaling regulates both the survival and proliferation of AT2 cells post-injury, as well as their migration and differentiation towards an AT1 phenotype (*14, 17*). Recent studies suggest that a subset of AT2 cells with activated Wnt-βcat-signaling display higher progenitor activity than the Wnt-inactive bulk population of AT2s (*11, 16, 18, 19*). Such differences in AT2 progenitor subsets are thought to be spatially induced in response to signals from the niche. For example, single cell RNA-sequencing data revealed that PDGFRα fibroblasts express Wnts, suggesting they might control the local activation of β-catenin AT2 cells (*19–21*), but the precise molecular signals and stromal cells that select AT2 progenitors are still undefined (*16, 19, 21*). Furthermore, whether and how these signals responsible for maintenance and repair of the AT2/mesenchymal niche are affected by hypercapnia and other signals in the injured lung injury are not known.

Here, we determined the effects of hypercapnia on AT2 progenitor capacity using unbiased RNA-sequencing analysis of flow-sorted AT2 cells and validation with lineage-labelled AT2 cells subjected to *ex vivo* organoid growth. Our findings suggest that hypercapnia limits AT2 cell β-catenin signaling and progenitor function by altering Wnt-expression in surrounding niche cells. We show hypercapnia skews expression of PDGFRα^+^/stroma-derived Wnt signals away from those typically known to activate β-catenin signaling (e.g., canonical Wnt2), and towards those historically shown to drive morphogenetic processes independently of β-catenin (e.g., non-canonical Wnt5a). We validate this model showing Wnt5a inhibits β-catenin signaling in primary AT2 *ex vivo* cultures. We also show *Pdgfra/Wnt2* cells are spatially closer to AT2 cells than *Pdgfra/Wnt5a* cells, and that PDGFRα^+^ cells express these two Wnts non-uniformly, suggesting that their spatial arrangement could direct proliferative versus differentiative zones in the distal lung. Altogether, these data suggest a mechanism by which hypercapnia may slow lung repair after injury, with broad implications for understanding how stromal-derived Wnt signals direct alveolar epithelial behaviors during repair.

## RESULTS

### Hypercapnia limits AT2 proliferation *in vitro* and *in vivo*

To assess the effect of hypercapnia on AT2 cell progenitor capacity, we employed a 3D-organoid model. Mouse primary AT2 cells (mAT2) were isolated by fluorescence-activated cell sorting (FACS) (EpCam^+^MHCII^+^) from leukocyte/endothelial cell-depleted lung homogenates of wild-type mice as described (*22, 23*) (**Fig. S1A),** or from *Sftp^CreERT2^:R26R^EYFP^* mice as EpCam^+^YFP^+^ (**Fig. S1B**). In *Sftp^CreERT2^:R26R^EYFP^* mice, tamoxifen administration permanently induces the expression of YFP specifically in AT2 cells and their progeny (*18*). To establish lung organoids, AT2 cells were embedded in Matrigel in the presence of mesenchymal cells in a 1:10 ratio **(Fig. 1A**). In normocapnic conditions (5% CO_2_, NC), sphere-like colony formation was seen typically between 5-7 days, which continued for up to 21 days (**Fig. S1C**). When cultures were started in the presence of high CO_2_ (20% CO_2_, HC), we did not observe organoid formation (data not shown), suggesting hypercapnia profoundly affects AT2 cell colony formation. To better assess consequences of hypercapnia for AT2 progenitor activity, we grew organoids for 7 days in control media before switching cultures to conditioned-media equilibrated to either 5 or 20% CO_2_ for an additional 7-14 days (**Fig. 1, A and B**). Under these conditions, hypercapnia reduced AT2 clonogenicity (Colony Forming Efficiency (CFE)) and significantly decreased average organoid diameter compared to organoids grown in normocapnia (**Fig 1, C and D** and **Fig. S1C)**. Whole organoids were fixed and stained using Surfactant protein C (SftpC) and Hop Homeobox (HOPX) as markers for AT2 and AT1 cells, respectively. In normocapnic conditions, organoids displayed a characteristic structure with central AT1 cells surrounded by peripheral AT2 cells, whereas during hypercapnia, few AT2 cells were observed with no AT1 cell marker expression (**Fig. 1E**). We next assessed effects of hypercapnia on proliferation *in vivo*. Mice were exposed for 21 days to either breathing room-air (RA) or 10% CO_2_ (HC). There were modest changes in overall lung histology (**Fig. S1D**), consistent with previous reports that hypercapnia negatively impacts alveolar epithelial function (*24, 25*). Hypercapnia limited the proliferative capacity of AT2 cells in mice, as evidenced by fewer SftpC^+^AT2 cells co-expressing the S-phase marker, Ki67 (**Fig 1, F and G**).

**Fig. 1.**
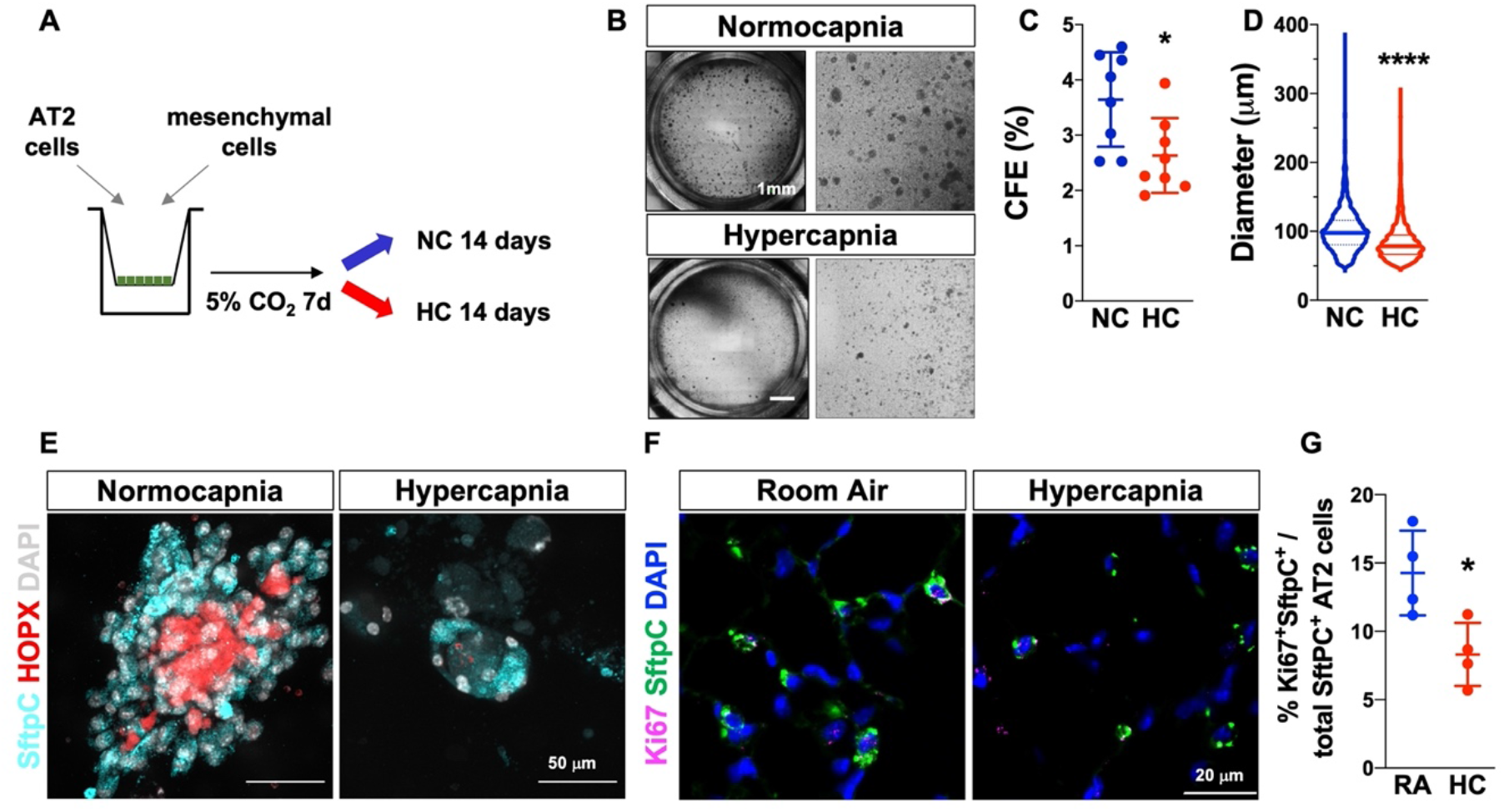
Hypercapnia limits AT2 cell proliferation in 3D culture organoids and *in vivo*. (**A**) Schematic of experiments designed to co-culture AT2 and mesenchymal cells in organoid assays, depicting the switch to normocapnia (5% CO_2_; NC) or hypercapnia (20% CO_2_; HC) media. (**B**) Left panels: Representative brightfield images of typical day 21 organoid cultures in normocapnia or hypercapnia. Right panels: Magnified views. (**C** and **D**) Graphs depict the inhibitory effect of hypercapnia on colony forming efficiency (CFE) and organoid size after 14 days in NC or HC. n=8. In (**D**) graph depict median with interquartile range. (**E**) Immunofluorescence analysis of SftpC^+^ (AT2) and HOPX^+^ (AT1) revealed a reduction in AT2 cells proliferation in organoids exposed to normocapnia or hypercapnia for 14 days. (**F** and **G**) Hypercapnia decreases the number of cells expressing Ki67 in the alveolar region of the adult mouse lung exposed to room air (RA) or HC for 21 days, as revealed by immunofluorescence. n=4. White arrows indicate SftpC^+^-Ki67^+^ AT2 cells. (C,D,G) Student’s t-test. \**P*<0.05; \*\*\*\**P*<0.0001.

### Hypercapnia inhibits Wnt-βcat signaling in AT2 cells

To determine how hypercapnia limits AT2 progenitor capacity and differentiation, we interrogated transcriptional differences in bulk-sorted AT2 cells isolated from mice exposed either to room air or hypercapnia. Mice were exposed to high CO_2_ for 7 and 21 days and AT2 cells were isolated from single cell suspensions as described above, RNA was isolated and bulk RNASeq was performed as previously described (*26, 27*). After 7 days of hypercapnia, only 15 differentially expressed genes (DEG) were identified compared to AT2 cells isolated from mice kept at room air (**Fig. 2, A and B**). After 21 days of exposure to hypercapnia, over 1200 DEG were identified compared to RA mice (**Fig. 2C**), where enrichment analysis of biological processes revealed hypercapnia inhibits lipid synthesis/metabolism, lysosomal pathways and alveolar growth (**Fig. 2D**). Specifically, hypercapnia limited the expression of genes coding for essential regulators of AT2 lineage (*Etv5, Abca3*) (*28, 29*), functional markers of AT2 cell function (*Sftpc, Nkx2.1*) (*28, 30*) and lipid metabolism (*Fabp5, Hmgcr*) (*28, 30*) (**Fig S2A-F),** suggesting an impairment of AT2 cell maturation and function. Decreased expression of SftpC was also confirmed by biochemical and immunological methods (**Fig. S2G-J**). Fibroblast growth factor receptor 2 (*Fgfr2*), which is necessary for AT2 cell maintenance and self-renewal (*31, 32*), was decreased by 50% in AT2 cells after 21 days of hypercapnia compared to room-air exposed mice (**Fig. S2K**), consistent with our evidence that hypercapnia attenuates AT2 proliferative capacity.

**Fig. 2.**
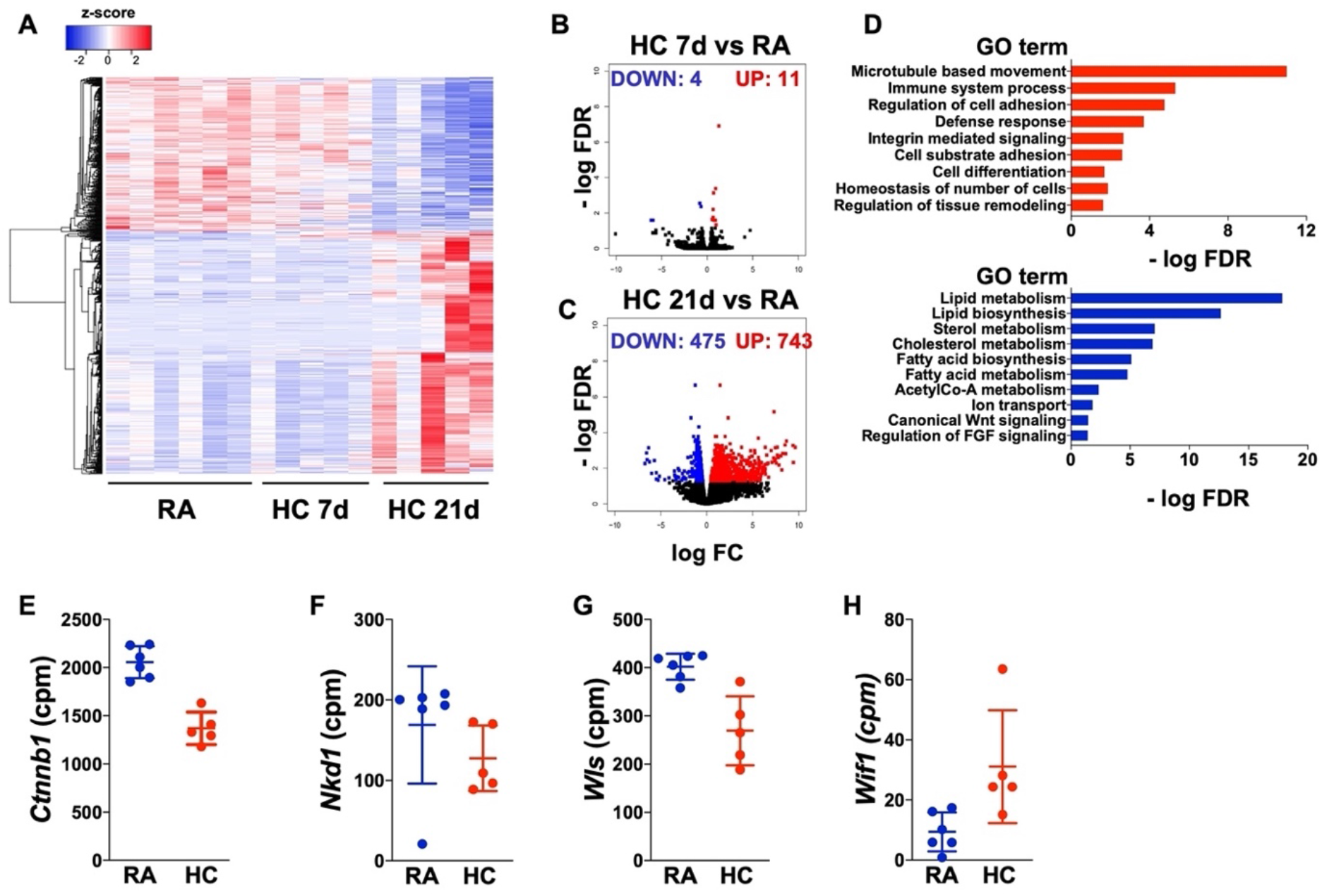
Transcriptomic analysis of isolated AT2 cells reveals inhibition of β-catenin signaling during hypercapnia. Bulk RNASeq was performed on flow cytometry sorted AT2 cells from mice breathing room air (RA, n=6) or exposed to 10% CO_2_ (HC). (**A**) Heatmap showing the clustering of differentially expressed genes (FDR *q* < 0.05) in AT2 cells after 7 (n=5) or 21 (n=5) days of hypercapnia exposure. (**B** and **C)** Volcano plots. (**D**) GO biological processes. (**E**-**H**) Expression of selected DEG (FDR *q*< 0.05) regulated by hypercapnia involved in the Wnt-βcat pathway.

Since Wnt signaling plays a major role in lung homeostasis and repair after injury (*18, 19, 33*), we interrogated established target genes and components related to this pathway. We found hypercapnia inhibited expression of *Ctnnb1* itself, as well as *Nkd1* (a known target) (*34*) and *Wls* (required for Wnts secretion) (*35*), and increased expression of *Wif1* (*36*), a negative feed-back regulator of Wnt-βcat signaling (**Fig. 2, E to H**). We confirmed this finding in AT2 cells isolated from mice exposed to hypercapnia for 21 days by following *Axin2* expression, the most widely expressed target of βcat signaling (*18, 19, 37*) (**Fig. 3A**). We also found hypercapnia limited the number of *Axin2*-expressing AT2 cells (*Sftpc*^+^*Axin2*^+^) using multiplexed in situ hybridization of wild-type mouse lung sections (**Fig. 3, B and C and Fig. S3A**), and independently confirmed this finding in the βcat signaling reporter mouse *Axin2^CreERT2-TdTom^* (**Fig. 3D**). In this Wnt-responsive fluorescent reporter line, *Axin2*^TdTomato+^ cells are restricted to the distal epithelium and surrounding mesenchyme at E13.5, and this pattern continues through adulthood (*18*). As with previous reports, we isolated bright populations of *Axin2*^TdTomato-HIGH^ expressing cells that are *SftpC* negative, as well as dim *Axin2*^TdTomato-DIM^ *SftpC*^+^ AT2 cells (**Fig. S3B-D**). Together, these data suggest that hypercapnia appears to antagonize AT2 progenitor activity by limiting βcat signaling within AT2 cells.

**Fig. 3.**
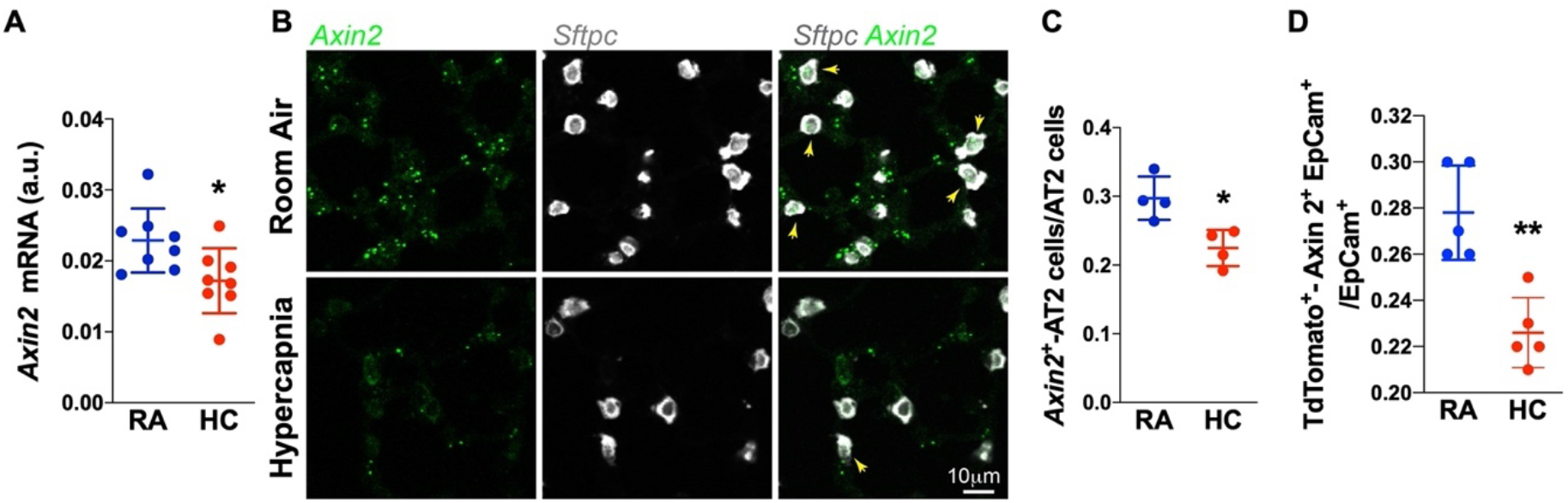
Hypercapnia decreases the number of Wnt-responsive AT2 cells. AT2 cells were isolated from mice exposed to room air (RA) or 10% CO_2_ (HC) for 21 days. (**A**) mRNA was isolated, and RT-qPCR was performed. n=8 mice. (**B and C**) *In situ* RNA hybridization showing decreased number of *Axin2*^+^-AT2 cells in mice exposed to hypercapnia. Yellow arrows indicate SftpC^+^-Axin2^+^ AT2 cells. n=4 mice. (**D**) Number of TdTom^+^Axin2^+^-AT2 cells from *Axin2^CreERT2-TdTom^* mice was determined by flow cytometry. n=5 mice. Graph shows the data from one of three independent experiments. \**p*<0.05; \*\**p*<0.01. Student’s *t-*test.

### Hypercapnia skews stromal-derived Wnts towards a βcat signaling inhibitory environment

To determine the mechanism by which hypercapnia limits βcat signaling activity in AT2 cells, we first interrogated our AT2 bulk RNAseq data set **(Fig. 2)** for altered expression of Wnt genes, given previous evidence that lung injury upregulates a set of Wnts in AT2 cells that could alter “bulk” AT2 behavior (*19*) and that short-term hypercapnia up-regulates Wnt7a in mouse lung homogenates (*38*). However, *Wnt7b*, which is highly expressed in AT2 cells (*19*) was not altered by hypercapnia (data not shown). While *Wnt3a* and *Wnt4* expression were mildly reduced by HC, they were expressed at very low levels and were not validated by RNAfish (not shown) or single cell data (*27*). Since stroma niche cells proximal to AT2 cells are critical for AT2 proliferation and differentiation to AT1 cells (*13, 16*), we sought to assess whether hypercapnia limits alveolar epithelial cell renewal by modifying Wnt signals produced from adjacent fibroblasts. We isolated PDGFRα-expressing fibroblasts (CD45^-^CD31^-^Epcam^-^PDGFRα^+^) from mice exposed to high CO_2_ for 10 days **(Fig. S4A)** and analyzed their transcriptomic profiles. We identified ~310 genes differentially expressed by exposure to hypercapnia **(Fig. 4A).** Analysis revealed robust expression of a number of Wnts previously identified in our scRNA-seq data sets as fibroblast-enriched (Wnts 2, 2b, 9a, 4, 5a and 11) **(Fig. 4B)** (*27*), where Wnt5a and Wnt 2 were the most abundant. Hypercapnia increased *Wnt5a* (significantly) a Wnt typically associated with inhibition of βcat signals and promoting cell shape changes (*39, 40*). Conversely, expression of Wnt2, which has been described to activate canonical Wnt signaling in rat lung fibroblasts and in NIH-3T3 (*41, 42*), trended downwards The hypercapnia-induced *Wnt5a* increase and Wnt2 decrease were independently confirmed by isolating mRNA from PDGFRα^+^ fibroblasts (**Fig. 4, C and D).** Multiplexed *in situ* hybridization (RNAfish) demonstrated that rather than increasing the number of *Wnt5a*^+^ cells, hypercapnia appeared to increase *Wnt5a* signal from cells already expressing this Wnt **(Fig. 4D and Fig. S4B**).

**Fig. 4.**
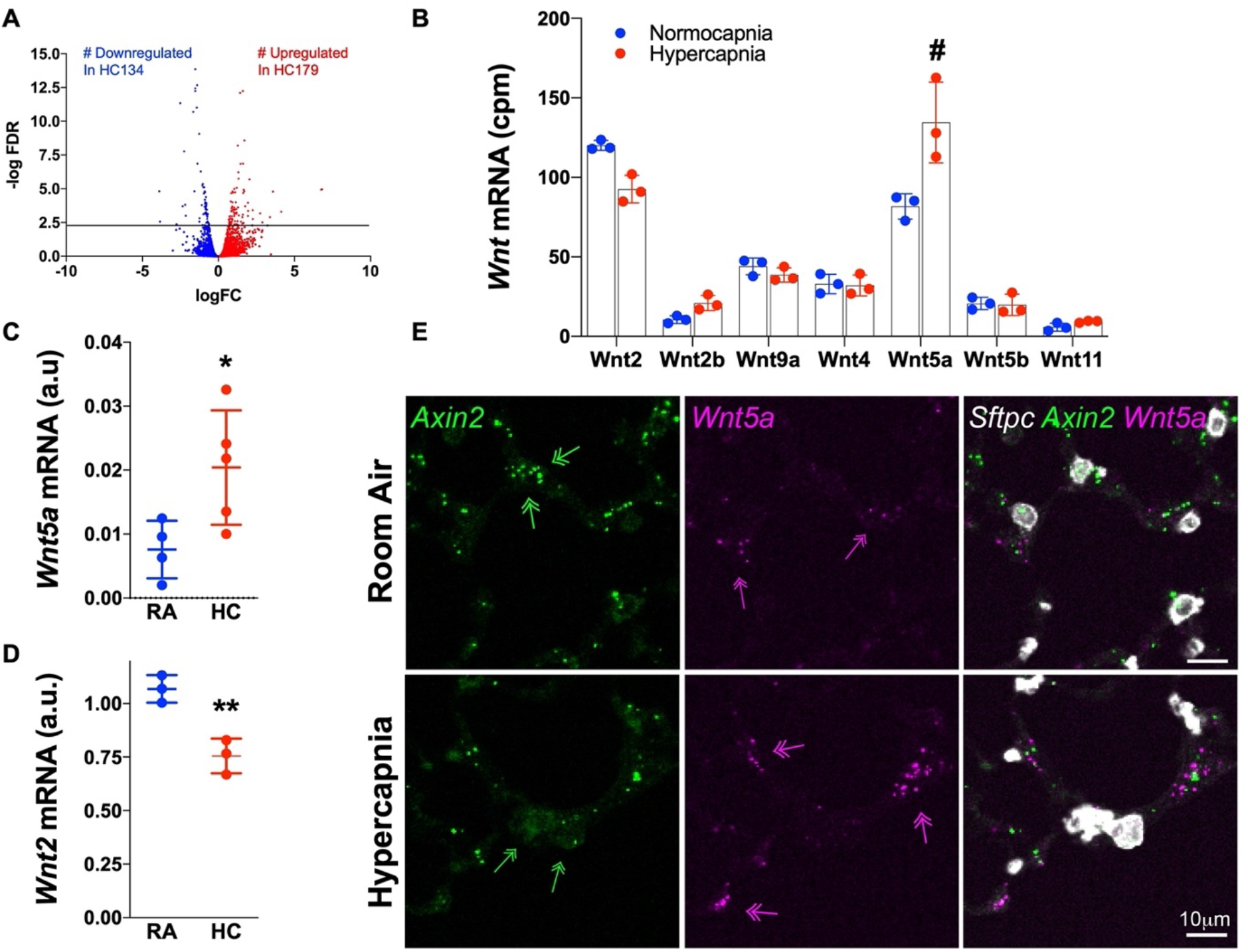
Hypercapnia increases Wnt5a expression in PDGFRα fibroblasts. (**A**) Lung PDGFRα fibroblasts were isolated via flow cytometry cell sorting from mice breathing room air or exposed to 10% CO_2_ (HC) for 10 days and analyzed using population RNA-Seq. n=3, with cells isolated from 3 mice in each replicate. (**A**) Volcano plot showing hypercapnia-down and up regulated genes. (**B**) Expression of Wnt genes in PDGFRα^+^ fibroblasts. # Indicates DEG (FDR q< 0.05). (**C** and **D**) mRNA was isolated, and RT-qPCR was performed. (**C**) Wnt5a (n=4) and (**D**) Wnt2 (n=3). (**E**) Representative image of in situ RNA hybridization showing increased expression of Wnt5a outside AT2 cells in mice exposed to hypercapnia. Green arrows indicate *Axin2* and magenta arrows indicate positive cells. n=4 mice. \**P*<0.05; \*\**P*<0.01. Student’s *t*-test.

Validating whether particular Wnts are βcat “activating” or “inhibiting” is important, given conflicting reports on the role of Wnt5a in AT2 progenitor behavior (*19, 39, 43*). As is typically observed, we found recombinant Wnt3a (rWnt3a) promoted βcat signaling in alveolar epithelial (A549) cells, whereas Wnt5a did not, using an established reporter assay **(Fig. 5A).** rWnt3a is used as a canonical Wnt/βcat activator (positive control), since validated recombinant Wnt2 (i.e., purified from mammalian cell culture secretions) is not available (R&D Systems; Bio-techne). We independently confirmed these findings by expressing Wnts via adenoviral transduction of primary AT2 cell cultures, showing Wnt5a limits, whereas Wnt3a elevates the βcat signaling pool, using an established affinity precipitation-based assay to capture and quantify the cadherin-free signaling pool of βcat (**Fig. 5, B and C**) (*44*). Together, these data show that Wnt5a antagonizes βcat signaling in cultured AT2 cells and raise the possibility that hypercapnia may limit the progenitor capacity of AT2 cells through stromal-derived Wnt5a.

**Fig. 5.**
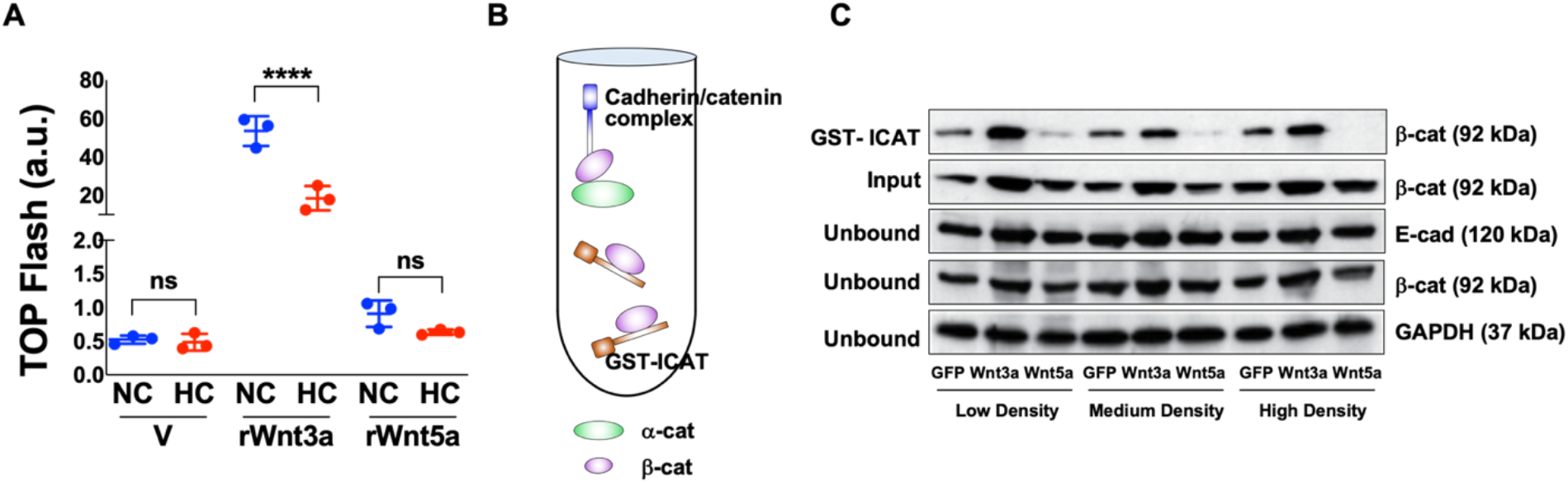
Wnt5a leads to a reduction in βcat signaling pool in alveolar epithelial cells. (**A**) β-Catenin-transcriptional activity was measured using the established, T cell-factor (TCF) Optimal luciferase-based Promoter (TOPFlash). A549 cells were transiently transfected with the reporter plasmid for 24 h, and then stimulated with recombinant Wnts for another 16 h. Cell lysates, normalized to equal protein concentrations, were assayed for luciferase activity. V=Vehicle. n=3 independent experiments run in duplicate. (**B**) Schematic for assay to quantify the cadherin-free signaling pool of β-Cat (β-Catenin) in cell lysates via GST-ICAT affinity precipitation as described in the Methods section. Inhibitor of Catenin and TCF (ICAT) is an 81 amino acid polypeptide that binds the central armadillo-repeat region of βcat and can be used to quantify the Wnt-stabilized pool of βcat. α-Cat (α-Catenin). (**C**) Immunoblot from rat AT2 cells plated at different densities and infected with adenoviruses coding for GFP, Wnt3a-IRES-GFP or Wnt5a-IRES-GFP and subjected to GST-ICAT affinity precipitation as in B. Input lysates and post-affinity precipitation (unbound lysates) are shown as controls. Note that across all cell plating conditions, Wnt3a increases whereas Wnt5a inhibits the GST-ICAT-bound (i.e., signal pool) of βcat. \*\*\*\**P*<0.0001; ns= not significant. One way ANOVA with Sidak’s post-comparison test.

Given recent evidence that the AT2 cell niche may comprise as few as 1-2 stromal cells to modulate βcat signaling in AT2 progenitor (*19, 21*) we sought to measure the spatial proximity of *Wnt5a* signal to *Sftpc^+^-AT2* cells compared with *Wnt2* using RNAfish. We focused on *Wnt2* for comparison with *Wnt5a* as it is one of the most abundant stromal-derived Wnts (*21, 27*) and is known to activate βcat signaling across diverse cell types (*45, 46*). Given that Wnt5a can antagonize the βcat signaling pool in AT2 cells (**Fig. 5),** we reasoned βcat-activating and - inhibiting Wnts might be spatially segregated in the AT2 niche to control AT2 progenitor versus differentiation decisions. By converting RNAfish signal to objects or spots based on signal intensity (see Methods section), we measured mean shortest distances between signal pairs **(Fig. 6, A and B).** We found that while both *Wnt2-* and *Wnt5a*-expressing *Pdgfrα* cells are spatially proximal to *Sftpc*^+^ AT2 cells (e.g., within ~8μm; **Fig. 6C**, the *Wnt2* signal is significantly closer than *Wnt5a* **(Fig. 6D**). Indeed, a histogram of the data show that 32% of the *Wnt2* signal is within 6μm of Sftpc signal, whereas only 19% of the *Wnt5a* signal is within this distance **(Fig. S5A)**. To assess whether *Pdgfra*^+^ cells comprising the AT2 progenitor niche are uniformly *Wnt2/Wnt5a*-double-positive, or instead capable of Wnt expression heterogeneity that could spatially-separate βcat-activating from βcat-inhibitory Wnts (e.g., *Wnt2* from *Wnt5a*, respectively), we isolated PDGFRα^+^ cells by flow cytometry and performed RNAfish analysis after cytospin **(Fig. S5B).** Interestingly, we found the PDGFRα^+^ population can be distinguished as 4-subtypes regarding *Wnt2/Wnt5a* expression: *Wnt2*^+^, *Wnt5a*^+^, *Wnt2/Wnt5a*^++^ and *Wnt2/Wnt5a*-negative **(Fig. 6E).** While PDGFRα^+^ cells no doubt express other Wnts, the results showed that this population is not uniformly positive for both *Wnt2* and *Wnt5a*, where these Wnts show different average distances from AT2 cells in lung sections, suggesting the possibility that PDGFRα^+^ stroma cells establish a spatial code of βcat-activating versus -inhibitory Wnt ligands to narrowly control the AT2 cell proliferative zone of the alveolus.

**Fig. 6.**
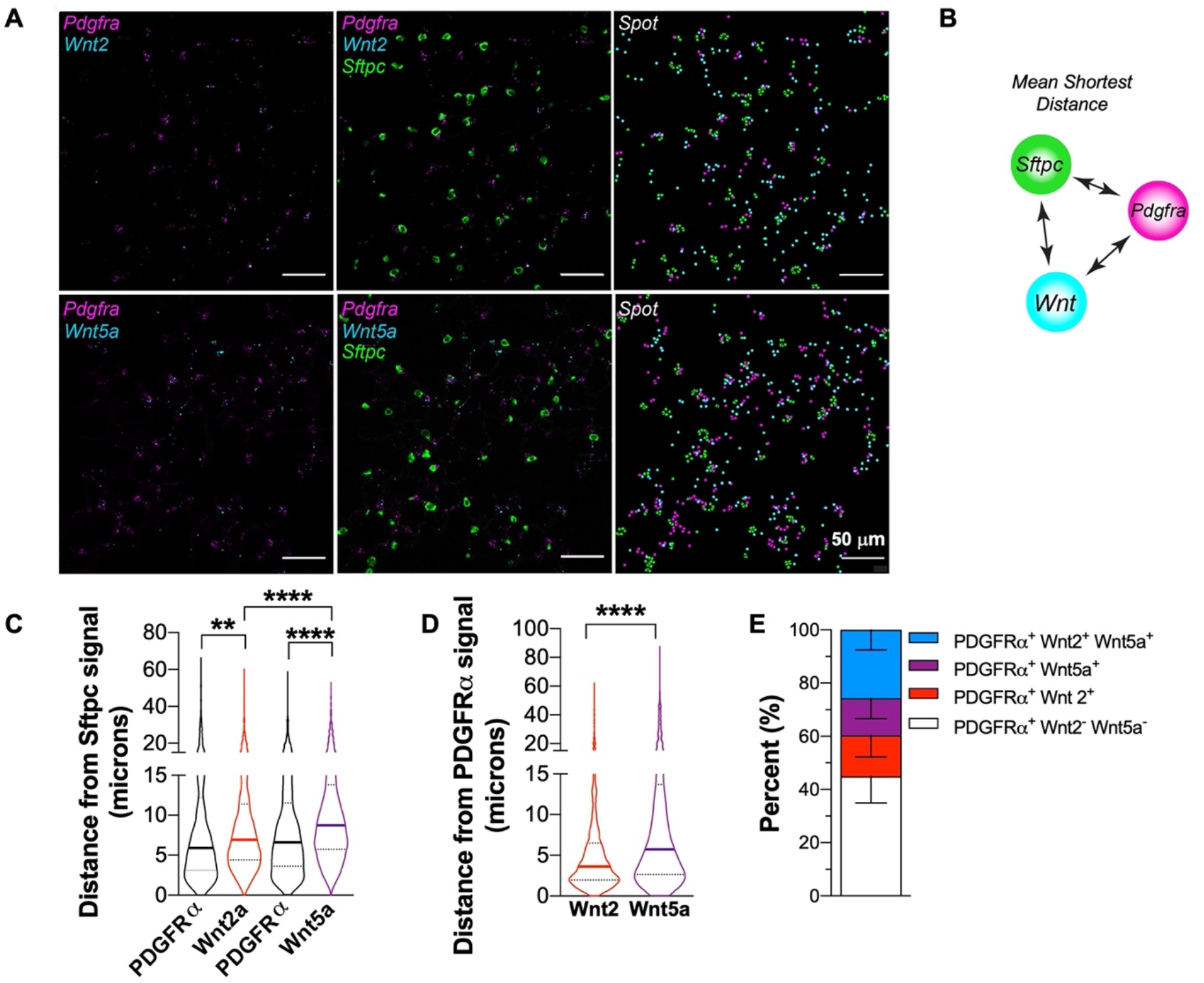
PDGFRα^+^ Wnt2 expressing fibroblasts are spatially closer to AT2 cells than Wnt5a-expressing fibroblasts. **(A**) Confocal images of *Pdgfra, Sftpc, Wnt2* and *Wnt5a* mRNA signal in lung tissue from wild-type mice. RNAFISH signal intensity converted to object spots to measure shortest distance between signals. (**B**) Schematic of spatial distance mapping algorithm. See methods for details. (**C** and **D**) Graphical representation of mean shortest distance of *Wnt2* or *Wnt5a* spots from *Sftpc* or *Pdgfra* signal, respectively. Graphs depict median with interquartile range. From 2 mice, 3 fields of view/mouse with more than 1500 measurements per condition. \*\**P*<0.01; *** *P*<0.001; \*\*\*\**P* <0.0001. One way ANOVA with Tukey’s post-comparison test. (**E**) Graph shows quantification of data from cytospin images of PDGFRα-flow sorted fibroblasts subjected to RNAFISH with probes to *Pdgfra, Wnt2* and *Wnt5a*. n= 3-independent PDGFRα-flow isolations pooled from 3 mice/each.

### Activation of βcat signaling rescues AT2 progenitor activity during hypercapnia

While hypercapnia can limit βcat signaling activity, the aggregate effects of hypercapnia are likely to be pleotropic. Thus, we sought to test whether the ability of hypercapnia to limit AT2 progenitor capacity could be offset by forced-activation of β-catenin signaling. The Wnt-β-catenin signaling pathway activators work by inhibiting glycogen synthase kinase-3β (GSK-3β), the central, inhibitory kinase in this pathway (*47*). GSK-3β inhibitors antagonize β-catenin phosphorylation and degradation, allowing β-catenin to then accumulate in the nucleus and propagate Wnt signals. As such, organoids were incubated in control media for 7 days and 24 h after switching the media to NC/HC they were incubated in the presence or absence of the GSK-3β inhibitor CHIR99021 (CHIR, 20nM). We observed that in normocapnia o hypercapnia conditions, the presence of CHIR increased the size of the organoids (**Fig. 7, A and B**). Remarkably, CHIR treatment rescued organoid growth during exposure to hypercapnia, although GSK3 inhibition also increased the size of organoids cultured in normocapnic conditions (**Fig. 7, A and B).** Since GSK3 plays roles in many signaling processes, we also assessed whether these effects are specific for βcat or other targets of GSK3 using AT2 cells isolated from *Sftpc^CreERT2^:Ctnnb1^Exon3fl/+^:R26R^EYFP^* mice (*48*). In these mice, Cre-dependent removal of a phosphodegron in exon3 generates a constitutively active form of βcat. Similar to the incubation with CHIR, the presence of constitutively active βcat rescues the effect of hypercapnia on organoid size (**Fig. 7, C and D).**

**Fig. 7.**
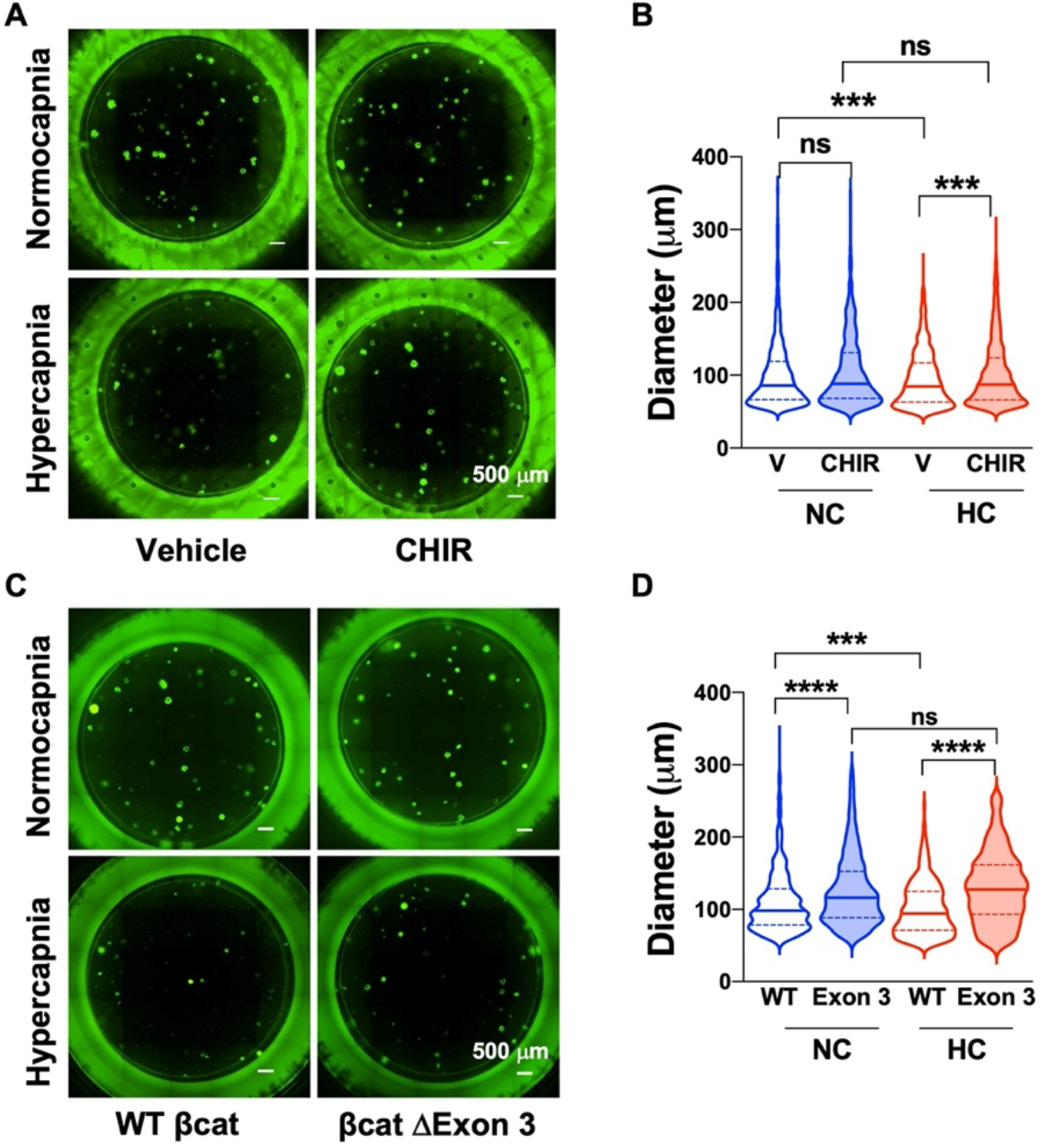
Activation of βcat signaling rescues AT2-proliferative capacity during hypercapnia. (**A**) Representative fluorescent images of typical day 21 organoid cultures. AT2 were isolated from *Sftp^CreERT2^:R26R^EYFP^* mice. Organoids were cultured in normocapnia (NC) or hypercapnia (HC) with or without the addition of CHIR as described in Fig. 1. (**B**) Graph depicts the effect CHIR treatment on NC and HC organoid size. n=8 mice in 4 independent experiments. (**C**) Representative fluorescent images of typical day 21 organoid cultures of AT2 isolated from *Sftp^CreERT2^:Ctnnb1^wt/wt^:R26R^EYFP^* mice (WT βcat) or *Sftp^CreERT2^:Ctnnb1^flExon3fl^:R26R^EYFP^* (βcat ΔExon 3). Organoids were cultured in NC or HC as described in Fig. 1. (**D**) Graph depicts the effect WT βcat vs βcat ΔExon 3 expression on NC and HC organoid size. n=4 mice of each strain in 2 independent experiments. (**B** and **D**) Data are shown as median with interquartile range. \**P*<0.05; \*\*\**P* <0.001; \*\*\*\**P* <0.0001; ns=not significant. One way ANOVA with Sidak’s post-comparison test.

## DISCUSSION

We describe a mechanism by which hypercapnia, an inevitable consequence of a lung protective ventilation strategy, slows or prevents lung repair after injury. We found that hypercapnia shifts the repertoire of Wnt signals in mesenchymal cells spatially localized in the niche of AT2 progenitor cells. Specifically, hypercapnia reduced expression of Wnt ligands that promote βcat signaling in AT2 cells and enhanced the expression of Wnt ligands that inhibit it. These changes were restricted to microdomains surrounding AT2 cells where they served to reduce AT2 proliferation and impair differentiation. Our results suggest that factors within the microenvironment of the injured lung, including hypercapnia, can disrupt signals from the mesenchyme required to restore the alveolar capillary barrier function and lung homeostasis after injury (**Fig. 8**).

**Fig. 8:**
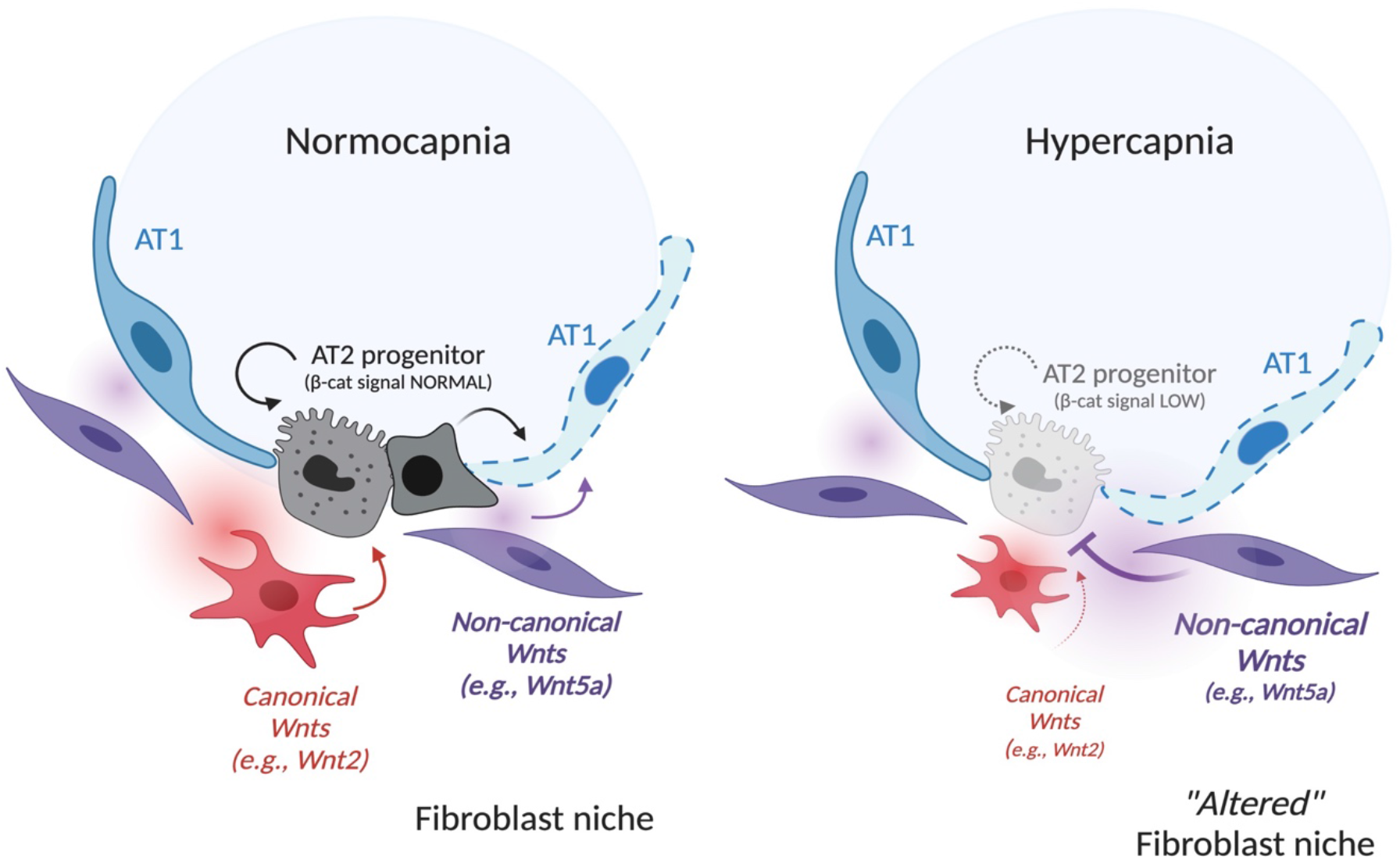
Model for hypercapnia-mediated inhibition of βcat signaling in AT2 cells via skewing of fibroblast-derived Wnts. **(Left**) Normocapnia: AT2 progenitors are spatially proximal to Pdgfrα/Wnt2-expressing fibroblasts (red/red gradient) that maintain βcat signaling and AT2 selfrenewal to replace damaged AT1 cells. Pdgfrα/Wnt5a*-*expressing fibroblasts (purple/purple gradient) are spatially further from the AT2 cell, perhaps to ensure separation of competing βcat-activating (Wnt2) from βcat-inhibiting (Wnt5a) signals. (**Right**). Hypercapnia leads to reduced βcat signaling in AT2 cells impairing cell renewal and differentiation by skewing Wnt expression in PDGFRα-stromal cells towards non-canonical variety, with *Wnt5a* significantly elevated. Narrowness of the AT2 progenitor niche raises possibility that elevated Wnt5a release (purple gradient) in close spatial vicinity to the Wnt2 signal (red gradient) antagonizes βcat signaling in AT2 cells, with inhibits proliferative capacity.

Previous lineage tracing experiments showed that a subgroup of AT2 cells, while retaining their surfactant biosynthetic capacity, can act as facultative progenitors (*19*). Our experiments suggest that the proliferative capacity of AT2 cells is decreased by hypercapnia. In a 3D-organoid model, the inhibitory effects of hypercapnia on the proliferative capacity of AT2 cells are so severe, that if the organoid incubation is started in hypercapnia equilibrated media, no organoids are formed. The decrease in AT2 proliferative capacity was also observed in lungs isolated from mice exposed to high CO_2_ for 21 days. To better understand the signaling pathways responsible for the hypercapnia-mediated decrease in proliferative activity, we analyzed the transcriptomic signature elicited in AT2 cells isolated from mice breathing room air or high CO_2_. We observed changes in the transcriptomic signature at 21 but not at 7 days of hypercapnia. At 21 days, hypercapnia decreased canonical AT2 markers, including *Sftpc* and genes involved in lipid metabolism (*Hmgcr, Fabp5*), as well as essential regulators of AT2-cell specification, such as *Etv5* and *Abca*. We also observed an increase in the expression of genes involved in processes of cell adhesion, tissue remodeling and cytoskeleton reorganization. Together, these data suggest hypercapnia causes a loss of AT2 cell characteristics.

Accumulating evidence suggest that canonical Wnt signaling pathway is essential for alveolar progenitor cells to induce alveolar epithelial repair (*18, 19, 33*). *Axin2* marks a sub-population of Wnt-responsive AT2 cells with progenitor characteristics (*16, 19*). Consistent with previous observations, we found that the Wnt activity (as evidenced by *Axin2* expression) is very low in control AT2 cells. A recent publication showed that Wnt5a expressed by fibroblasts in close proximity to AT2 cells induces canonical Wnt signaling and stimulates repair (*19*). Our data challenge the universality of this model, and instead reinforce a model where increased expression of *Wnt5a* by PDGRa-fibroblasts limits βcat signaling in AT2 cells. We validated Wnt-subtype functionality for a subset of these Wnts in primary AT2 cultures, showing that Wnt5a limits, whereas Wnt3a enhances βcat protein levels and signaling. Our results align with a recent lung organoid study showing that Wnt5a and Wnt5b ligands inhibit alveolar epithelial stem/progenitor expansion by impairing canonical Wnt signaling (*40*), as well as longstanding evidence that Wnt5a limits bcat signals in lung and other tissues (*49, 50*). Our data are also consistent with evidence that the mesenchymal population that sustains the self-renewal and differentiation of AT2 stem cells is positive for *Axin2, Pdgfra, Wnt2, Il6* and *Fgf7* (*21*).

Our results highlight the importance of experimental methods that provide single cell spatial resolution in studies of lung repair. Spatial transcriptomic approaches allowed us to localize these signals specifically to mesenchymal cells near AT2 cells. We found that freshly isolated PDGFRα^+^ cells can be distinguished as 4-subtypes regarding *Wnt2/Wnt5a* expression: *Wnt2*^+^, *Wnt5a*^+^, *Wnt2/Wnt5a*^++^ and *Wnt2/Wnt5a*-negative. These data suggest AT2 niche cells (i.e., PDGFRα stromal cells) may be targets of pathological conditions like hypercapnia. These data also suggest that the spatial arrangement of distinct PDGFRα^+^ stroma cell-subtypes may direct AT2 cell proliferative versus differentiative zones of the alveolus. Using RNAfish analysis, we found that *Wnt2*-expressing PDGFRα cells are spatially closer to *Sftpc^+^AT2* cells than *Wnt5*a-expressing PDGFRα cells. Together these data suggest that a more systematic evaluation of the major, stromal derived Wnts may shed light on how individual AT2 cells are selected for activation.

In summary, we demonstrate a fundamental effect of elevated CO_2_ levels on alveolar epithelial cell behavior which is regulated by Wnt signal modulation which inhibits the progenitor reparative AT2 cell function. These findings are of biological and clinical relevance as they pertain to patients with severe COVID-19 ARDS requiring mechanical ventilation. We contend that the hypercapnia-mediated mechanisms we uncovered are of relevance to lung repair after injury and thus contribute to the worse outcomes of these cohorts of patients if they are hypercapnic. Inhibition of AT2 proliferation in hypercapnic patients prevents the sealing of the epithelial barrier, increasing lung flooding, ventilator dependency and mortality.

## MATERIALS AND METHODS

### Mouse strains and Cre recombinase induction

Animal work was conducted in accordance with the recommendations in the Guide for the Care and Use of Laboratory Animals and the National Institute of Health. All procedures were approved by the Northwestern University’s Institutional Animal Care and Use Committee (Chicago, IL, USA; Animal Study Number: IS00010662). All strains including wildtype mice are bred and housed at a barrier- and pathogen-free facility at the Center for Comparative Medicine at Northwestern University. Animal experiments were performed on both male and female animals in all conditions, and animals were chosen at random from the cohort but not formally randomized. Adult (8 to 10 weeks old) C57BL/6J mice were obtained from the Jackson Laboratories (Strain Number: 000664) and were used as the wild type strain. *Sftpc^CreERT2^:R26R^EYFP^* and *Axin2^CreERT2-TdTom^,Ctnnbl^fl(ex3)/+^* (were kindly provided by Dr. Edward E. Morrisey, University of Pennsylvania) and their genotyping and characterization has been previously described (*18, 48, 51*).

For induction of estrogen-inducible Cre recombinase (Cre-ERT2) for conditional-tissue-specific conditional alleles *in vivo*, tamoxifen was dissolved in sterile corn oil (Millipore Sigma (T5648)) at 20 mg/ml concentration. Mice were injected intraperitoneally three times over the course of five days with 0.25 mg/g body weight to induce Cre-recombination of floxed alleles for lineage tracing in *Sftpc^creERT2^:R26R^EYFP^*(*18*).

Mice were provided with food and water ad libitum, maintained on a 14-hour light/10-hour dark cycle. For high CO_2_ exposure, mice were maintained in a BioSpherix C-Shuttle Glove Box (BioSpherix) for up to 21 days as described previously (*52*). Control mice were maintained in the adjacent space under room air conditions. At the selected time points, mice were euthanized with Euthasol (pentobarbital sodium–phenytoin sodium) and the lungs were harvested. We have previously shown that mice exposed to high CO_2_ had elevated PaCO_2_ and higher bicarbonate values after 3 days of exposure reflecting renal compensation of the respiratory acidosis (*53, 54*).

### Mouse AT2 cells isolation by flow cytometry

Tissue preparation for mouse AT2 isolation was performed as described (*22, 23*), with modifications. Multicolor flow cytometry and cell sorting were performed with an LSR Fortessa or BD FACSAria cell sorter using BD FACSDIVA software (BD Biosciences). Briefly, perfused lungs were treated with 50 U/ml dispase (Corning # 47743-724) and 0.25 mg/ml DNase (Sigma-Aldrich # D4513-1VL) and were subjected to manual dissection gently tearing and mincing the lung pieces. When required, single cell suspensions were enriched for epithelial cells using anti-EpCAM magnetic microbeads (Milteny Biotec # 130-105-958).

Lungs from wild-type, *Sftpc^CreERT2^:R26R^EYFP^* and *Axin2^CreERT2-TdTomato^* mice were processed into single-cell suspensions as indicated above. <u>Wild-type AT2 cells</u>: AT2 cells were sorted from the single-cell suspensions using antibody staining for CD31-PECy7 (eBioscience; #25-0311-81), CD45-PECy7 (eBioscience # 25-0451-82), and EpCAM-APC (eBioscience; # 17-5791-80). <u>Wildtype AT2 cells</u> CD45^-^CD31^-^EpCAM^+^ cells were gated for MHCII (eBioscience #48-5321-82) and selected as EpCAM^Int^ MHCII^high^ as previously described (*55*) and showed in **Fig. S1**. <u>SftpC (YFP)^+^-AT2 cells</u>: following negative selection for CD31 and CD45, YFP^+^-AT2 cells were isolated based on enhanced YFP (EYFP) fluorescence and sorted as indicated above and in **Fig. S1**. Axin2^+^-AT2 cells: following negative selection for CD31 and CD45, Axin2^+^-AT2 cells were positively gated for TdTomato and EpCAM as indicated above and in **Fig. S3**. Compensation, analysis, and visualization of the flow cytometric data were performed using FlowJo software (FlowJo,L.L.C., 10.7.1). “Fluorescence minus one” controls were used to set up gates.

### Alveolar Organoids

Clonal alveolar organoid assays were performed as described previously (*13, 18*). In brief, a mixture of AT2 cells (5 x10^3^ AT2, YFP^+^-AT2 or YFP^+^-βcat ΔExon 3 and lung fibroblasts (5 x 10^4^) in growth media containing the following: αMEM media (Thermo Fisher #41061029) supplemented with 4.5 g/l D-glucose, 2mM L-glutamine, 10% FBS and 1% penicillin-streptomycin, 1% insulin/transferrin/selenium (Thermo Fisher #41400045), 0.002% Heparin, 0.25 ug/ml Amphotericin B (Millipore Sigma #A2942), 2.5 μg/ml rock inhibitor Y24632 (Selleckchem #S1049) was used for the assays. Lung fibroblasts for organoids assays were isolated from adult wild type mice and plated in DMEM supplemented with 4.5 g/L D-glucose, 2mM L-glutamine, 10% FBS and 1% penicillin-streptomycin as previously described (*16*). Immediately before use cells were treated with mitomycin-C (Millipore Sigma #M4287) for 2 hours. Cells were then suspended in a 1:1 mixture of organoid growth media and growth factor-reduced, phenol-free Matrigel (Corning #356231). 100μl of the cell/media/matrigel mixture was then plated into individual 24 well cell culture inserts (Corning, #3470) and allowed to solidify at 37°C for 5 minutes. Organoid growth media was added into the bottom of the 24-well plate. After 5-7 days of culture, organoids were moved to normocapnic and hypercapnic conditions and grown for an additional 7-14 days. The growth media was buffered with an F-12/Tris/Mops solution with a 3:1:0.5 ratio as previously described (*8*) to maintain a pH of 7.35-7.45. CHIR was used at a concentration of 10 nM starting 24 hours after switching media. The media with or without treatment was changed every 48 hours. Organoids were imaged on a Nikon Ti^2^ Widefield in brightfield and GFP channels. In some cases at the end of the experiment organoids were fixed in 4% paraformaldehyde. Images were processed in Nikon Elements (5.11.00) and quantified using ImageJ/Fiji software (v152p) for organoid diameter. Alveolar organoids are defined as a clonal colony with a minimum diameter of 50 microns. The following antibodies were used HOPX (Santa Cruz Biotechnology; mouse monoclonal antibody; #sc-398703, 1:50), SftpC (Millipore; rabbit polyclonal antibody AB37086; 1:250) and imaging was performed in a Nikon Eclipse Ti confocal microscope.

### Immunostaining of whole lung sections

After lungs were cleared of blood by perfusion with cold PBS via the right ventricle, they were inflated with 4% paraformaldehyde and allowed to fix overnight. Tissue was then dehydrated, paraffin embedded, and sectioned. Hematoxylin and eosin staining was performed to examine morphology. Immunohistochemistry was used to detect protein expression using the following antibodies on paraffin sections: SftpC (rabbit, Millipore, AB37086, 1:250), and Ki67 (eBioscience, SolA15 rat monoclonal antibody; #14-5698-8, 1:100). Following PBS washes, fluorophore-conjugated secondary antibodies were prepared at 1:500 for 60 minutes in the dark at room temperature. Hoechst was diluted 1:1,000 and applied for 15 minutes at room temperature following secondary antibody incubation. Coverslips and filters were mounted using ProLong Gold anti-fade solution. Images were obtained using Zeiss Axioplan epifluorescence. Cell counts were performed using ImageJ software.

### Isolation of mouse lung fibroblasts

Mesenchymal populations from whole lung were isolated from wildtype C57Bl/6 mice. Briefly, perfused lungs were inflated with digestion buffer containing 200 U/ml Collagenase (Millipore Sigma; Cat # C0130-500 mg); 4 U/ml Elastase (Worthington; Cat # LS002292), 0.25 mg/mL DNase (Sigma; Cat # D4513-1VL) and coarsely minced before processing in C-tubes (Miltenyi Biotec) with a GentleMACS dissociator (Miltenyi Biotec) according to the manufacturer’s instructions. Homogenate was passed through 40-μm nylon filter to obtain a single-cell suspension and subjected to red blood cell lysis reagent (BD Pharm Lyse, BD Biosciences). Cells were subjected to negative selection using CD45 magnetic microbeads (Miltenyi Biotec # 130-052-301). Fibroblasts cells were identified as CD45^-^ CD31^-^ EpCAM^-^ PDGFR^+^ by flow cytometry as outlined in **Fig. S4** using PDGFRA antibody (eBioscience #12-1401-81).

### Transcriptomic profiling via RNA-Seq

FACS-based isolation of AT2 cells or PDGFRα^+^ fibroblasts from wild type mice was performed at the indicated time points for each experiment as described above. The RNeasy Plus Micro Kit (Qiagen, 74034) was used to isolate RNA and remove genomic DNA. RNA quality was assessed with the 4200 TapeStation System (Agilent Technologies). Samples with an RNA integrity number (RIN) lower than 7 were discarded. RNA-Seq libraries were prepared from 5 ng total RNA using the NEB Next RNA Ultra Kit (QIAGEN) with poly(A) enrichment. Libraries were quantified and assessed using the Qubit Fluorimeter (Invitrogen, Thermo Fisher Scientific) and the Agilent TapeStation 4200. Libraries were sequenced on the NextSeq 500 instrument (Illumina) at 75 bp length, single-end reads. The average reading depth across all experiments exceeded 6 × 10^6^ per sample, and over 94% of the reads had a Q score above 30. Bioinformatics work was performed on “Genomics Nodes” using Northwestern High-Performance Computing Cluster, Quest (Northwestern IT and Research computing). Reads were demultiplexed using bcl2fastc (version 2.17.1.14). Quality was controlled with FastQC. Samples that did not pass half of the 12 assessed quality control (QC) statistics were eliminated. Then reads were trimmed and aligned to mm10 reference genome using TopHat2 aligner (version 2.1.0). Reads counts were associated with genes using the GenomicRanges (*56*). The downstream differential gene expression was conducted with edgeR R/Bioconductor packages (*57, 58*). Genes with less than 1 normalized counts from more than 3 samples were excluded from the analysis. Hierarchical clustering heatmaps were also performed for top 1,000 most differential expressed genes. Statistical significance was controlled at a *p*-value of 0.05 with False Discovery Rate (FDR) adjusting for multi-pairwise comparison. GO analysis was performed using GOrilla (*59*) on 2 unranked gene lists.

### Quantitative real time PCR

RNA was isolated from primary AT2 cells or PDGFRα^+^ fibroblasts and purified using RNeasy Plus Micro Kit before cDNA preparation with qScript cDNA synthesis kit (Quanta Bio, 95047). For qRT-PCR, IQ SybrGreen master mixes (Bio-Rad, 1708880) were used according to manufacturer’s instructions. The following primers were used:

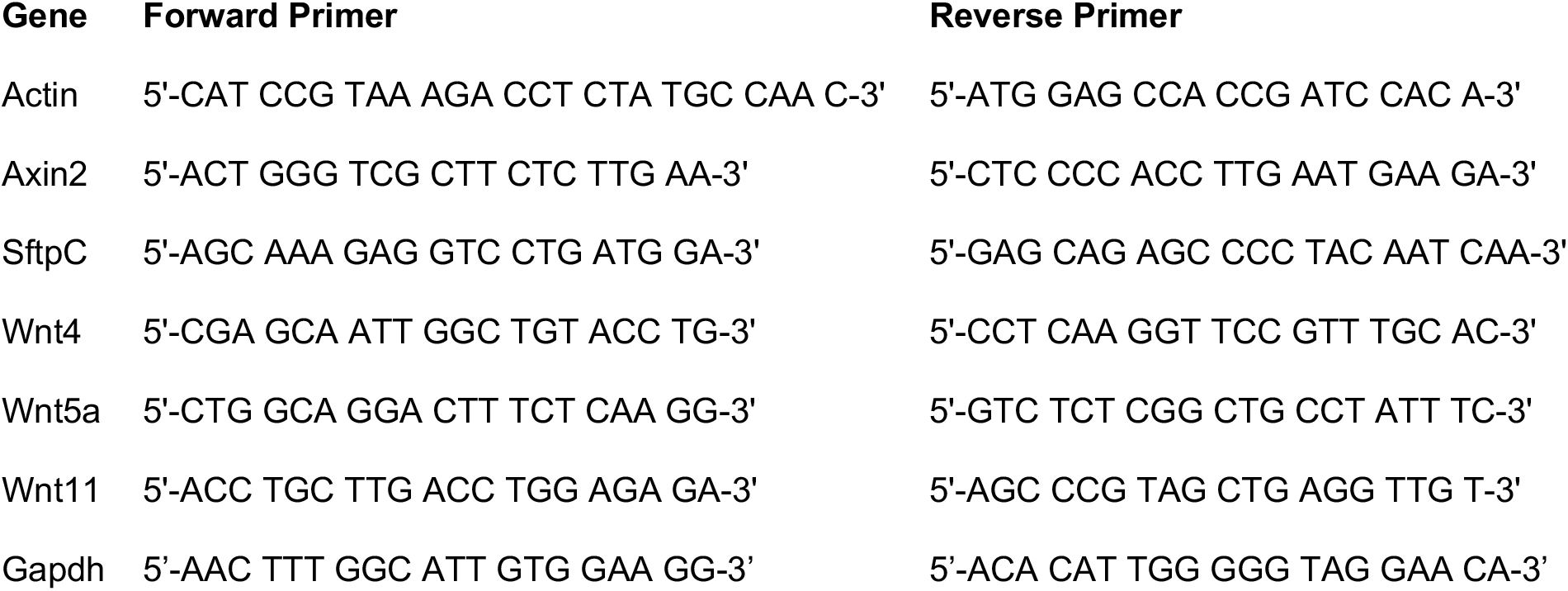

TaqMan qPCR was performed for Wnt2 using pre-designed TaqMan primer/probe sets (Life Technologies, Assay ID: Mm00470018 or Mm01156901) using Gadpdh (Assay ID: Mm99999915_g1) following manufacturer’s instruction. qPCR was performed in triplicate for each biological sample and fold enrichment was calculated from Ct values for each gene of interest, normalized to expression of housekeeping genes.

### Immunoblotting

Protein content in cell lysates was quantified by Bradford assay. Equal amount of proteins was resolved on polyacrylamide gels (SDS-PAGE). Proteins were transferred to a nitrocellulose membrane using a Trans-Blot-Turbo Transfer System (Bio-Rad). Incubation with specific primary antibodies was performed overnight at 4C. SftpC was detected using Millipore; rabbit polyclonal antibody AB37086 (1:2500) and actin using a rabbit polyclonal antibody from Sigma-Aldrich (#A2066; 1:8000). HRP-tagged secondary antibodies (Bio-Rad, 1721011 and 1721019) were used in combination with Super-Signal ECL kit (Thermo Fisher) to develop blots using a LI-COR Odyssey Imager and companion software Image Studio version 5.2. Blots were quantified by densitometry (ImageJ 1.46r; National Institutes of Health, Bethesda, MD).

### Single Molecule mRNA Fluorescent in Situ Hybridization (RNA-FISH)

Multiplex fluorescence in situ hybridization was performed using RNAscope (Advanced Cell Diagnostics, Newark, CA, USA). Mouse lungs were inflated to 15 cm H_2_O and fixed with 4% paraformaldehyde (EMS, Hatfield, PA, USA) for 24 h. Lungs were paraffin embedded and 5 μm tissue sections were mounted on Superfrost Plus slides (Thermo Fisher Scientific, Waltham, MA, USA). Slides were baked for 1 h at 60°C, deparaffinized in xylene and dehydrated in 100% ethanol. Sections were treated with H_2_O_2_ (Advanced Cell Diagnostics) for 10 min at room temperature and then heated to mild boil (98–102°C) in 1× target retrieval reagent buffer (Advanced Cell Diagnostics) for 15 min. Protease Plus (Advanced Cell Diagnostics) was applied to sections for 30 min at 40°C in a HybEZ Oven (Advanced Cell Diagnostics). Hybridization with target probes, pre-amplifier, amplifier, fluorescent labels and wash buffer (Advanced Cell Diagnostics) was done according to Advanced Cell Diagnostics’ instructions for Multiplex Fluorescent Reagent Kit version 2 and 4-Plex Ancillary Kit version 2 when needed. Parallel sections were incubated with Advanced Cell Diagnostics positive and negative control probes. Sections were covered with ProLong Diamond Antifade Mountant (Invitrogen). Mouse probes used were: *Wnt11*-C1 (#405021-C1); *Wnt4*-C2 (#401101-C2); *Wnt5a*-C2 (#316791-C2); *Sftpc*-C3 (#314101-C3); *Axin2*-C1 (#400331-C1), Mouse Negative Control #320871 all from Advanced Cell Diagnostics; Newark, CA. Opal fluorophores (Opal 520 #FP1487001KT; Opal 620 #FP1495001KT; Opal 690 #FP1497001KT) from Akoya (Perkin Elmer, Shelton, CT) were used at 1:1500 dilution in Multiplex TSA buffer (Advanced Cell Diagnostics). GSE164733 Images were captured on a Nikon A1R confocal microscope (Tokyo, Japan) with ×20 and ×40 objectives, followed by uniform processing and pseudo-colouring in Fiji (https://imagej.net/Fiji). To assess spatial proximity of *Wnt2*, *Wnt5a* and *Pdgfra* mRNA signals to *Sftpc*-positive AT2 cells, quantification was performed using Nikon NIS-Elements software version 5.11.01 using the General Analysis module, NearstObjDist algorithm. This converts the RNAfish signal into objects (circle/spot) based on signal density, so distances can be measured between spot centers. The smallest distance a given “spot” of channel/color (e.g., *Pdgfra*^+^ spot) is from the other two channel/color “spots” (e.g., *Sftpc*^+^ spot or *Wnt2*^+^ spot) is measured for all image spots. Mean shortest distance measurements were quantified and plotted in Prism.

### β-catenin/TCF Reporter Assay

A549 cells (1.5×10^5^ cell/well) were transfected with Super8-TOPflash (containing 8 TCF-consensus binding sites) or 1 μg of Super8-FOPflash (8 mutant TCF-binding sites) reporter plasmids (kindly provided by Randall Moon, University of Washington, Seattle) using Lipofectamine 2000 (Invitrogen); a thymidine kinase-Renilla plasmid (0.1 μg) was also included to normalize luciferase values to the efficiency of transfection as we previously described (*17*). After 4 hours of transfection, media was replaced, and cells were incubated overnight under normocapnic or hypercapnic conditions. The next morning rWnt3 (50ng/mL) or rWnt5a (50ng/mL) was added to the media and cells were incubated for another 24 hours. At the end of the incubation, A549 cells were solubilized using the Dual-Luciferase Assay Kit (Promega), and luciferase activity was quantified using a microplate dual-injector luminometer (Veritas) according to manufacturer’s instructions. Briefly, cells in each well of a 6 well plate were incubated on ice with 250uL of a 1x passive lysis buffer for 15 minutes followed by scrapping to lift adherent cells. 20uL of cell lysate was mixed with 100uL of supplied Luciferase Assay Reagent II and firefly luciferase was measured.

### Measurement of β-catenin signaling pool

AT2 cells were isolated from rat lungs as described (*17, 60*). 3×10^6^ AT2 cells were plated to 100mm (low density), 60mm (medium density), and 35mm (high density) culture dishes and immediately infected with 20 pfu/cell adenovirus (Vector Biolabs, 10^10^–10^12^ pfu/ml).): Ad-CMV-GFP, Ad-Wnt3a, and Ad-Wnt5a. Infection efficiency of (>90%) was confirmed by visualizing GFP expression in living cells. Cells were lysed after 72 hours as described (*17*). Protein lysates were processed for GST-ICAT pulldown as described (*44*). Briefly, 100 μg of lysate was saved as input fraction. 1000 μg of lysate was combined with 50ul of a 50:50 slurry of pre-washed glutathione-sepharose beads (GE 4B), and 20 μg of GST-ICAT and tumbled at 4°C for 2 hours. Samples were centrifuged at 10,000 rpm for 1 minute to pellet beads, and a 5 μl aliquot of supernatant was collected as the unbound fraction. Beads were washed with 0.1% triton buffer and centrifuged three times, and the final pellet resuspended in 35 μl 2X SDS loading buffer as bound pulldown fraction. Protein lysates were run on 6% acrylamide gel and transferred to nitrocellulose for immunoblotting as described above. Primary antibodies: mouse anti-ABC (Millipore Clone 8E7 #05-665) at 1:1000, mouse anti-β-cat (BD Clone 14 #610154) at 1:1000 or 1:5000, mouse anti-E-cadherin (BD Clone 36 #610182) at 1:1000 and rabbit-anti-GAPDH (Santa Cruz #sc-25778) at 1:1000.

### Statistics

Analyses of significance were performed using GraphPad Prism (v.8.4.2) software. A *p* value ≤0.05 was considered statistically significant. A standard two-tailed unpaired Student’s t-test was used for statistical analysis of two groups. One-way ANOVA, followed by analysis-specific posttests, was carried out when more than two variables were compared. Unless stated otherwise data are presented mean ± S.D. overlaid with individual data points representing replicates.

## Acknowledgements

The authors would like to thank Yair Romero Lopez, Fei Chen and Ziyan Lu for their technical support.

## Authors contributions

LAD, LCW, CJG, and JIS conceived and designed the research. LAD, LCW, NDM, PLB, DC, ASF, AW, MMH, HAV, MS, SMCM, CER performed and analyzed the experiments. LAD, LCW, NDM, PLB, AVM and CJG analyzed the experimental data. ZR and AVM performed the bioinformatic analysis. LAD, GRSB, CJG, JIS wrote the manuscript. All authors provided edits and feedback on the manuscript.

## Data and materials availability

All data needed to evaluate the conclusions in the paper are present in the paper and/or the Supplementary Materials. The RNA-Seq data sets are available at in the NCBI’s Gene Expression Omnibus (GEO) database #GSE139426 Room Air/Hypercapnia AT2 cells; for PDGFRα^+^-fibroblasts Room Air/Hypercapnia the record is process.

---

**Fig. S1.**
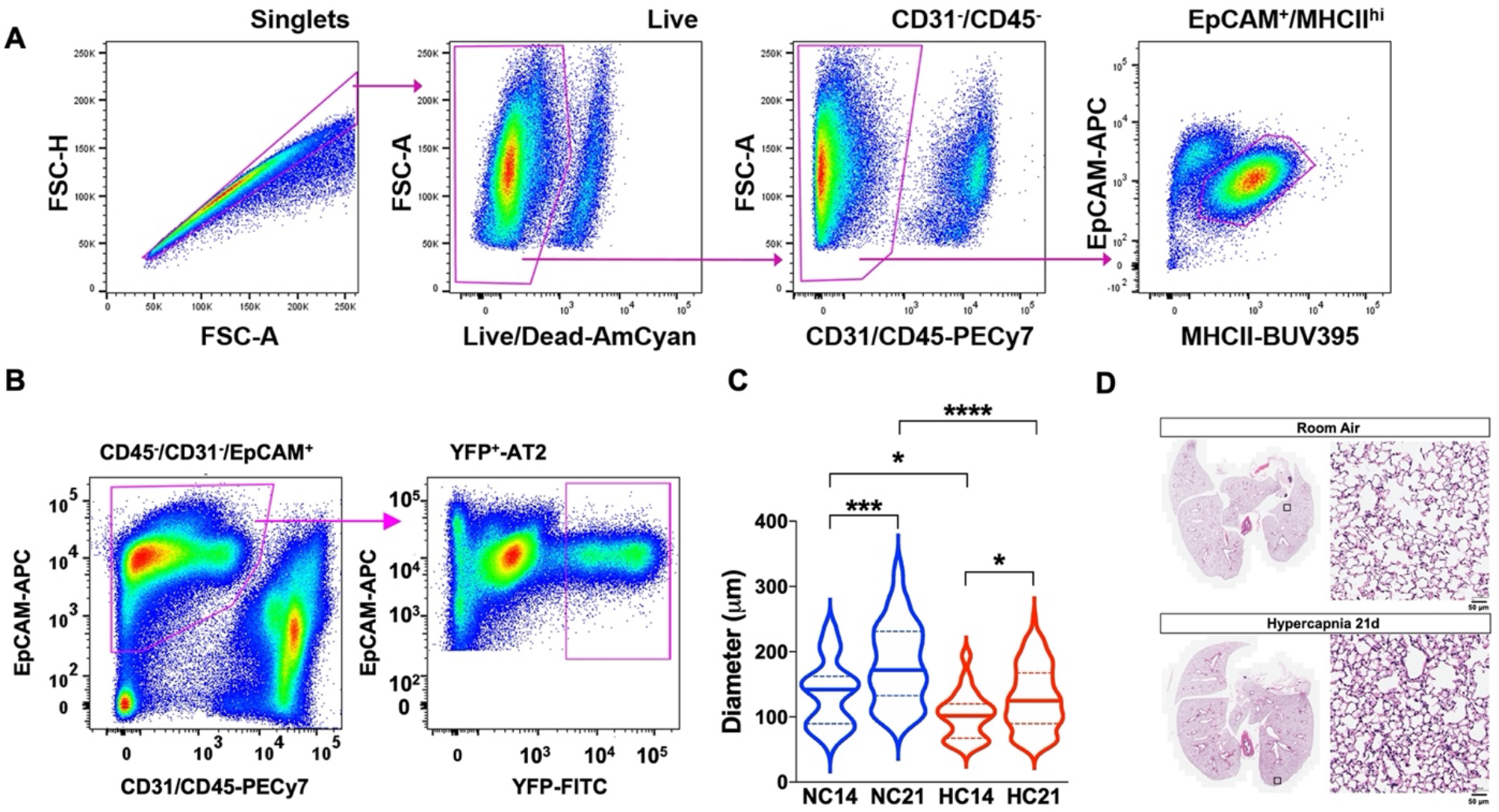
Hypercapnia limits AT2 cell proliferation in 3D culture organoids and *in vivo*. Gating strategies used to identify AT2 cells in lung single cell populations. Representative flow cytometry plots from a naïve adult mouse (**A**) and tamoxifen-treated *Sftp^CreERT2^:R26R^EYFP^* mice (B) are depicted. (**C**) Inhibitory effect of hypercapnia on organoids diameter after 7 and 14 days of exposure. Graph depicts median with interquartile range. \**P*<0.05; \*\*\**P*<0.001; \*\*\*\**P*<0.0001. One way ANOVA with Sidak’s post-comparison test. (**D**) Representative H&E of lung tissue from mice exposed to room air (RA) or 10% CO_2_ (HC) for 21 days showed that HC causes mild structural differences compared to RA.

**Fig. S2.**
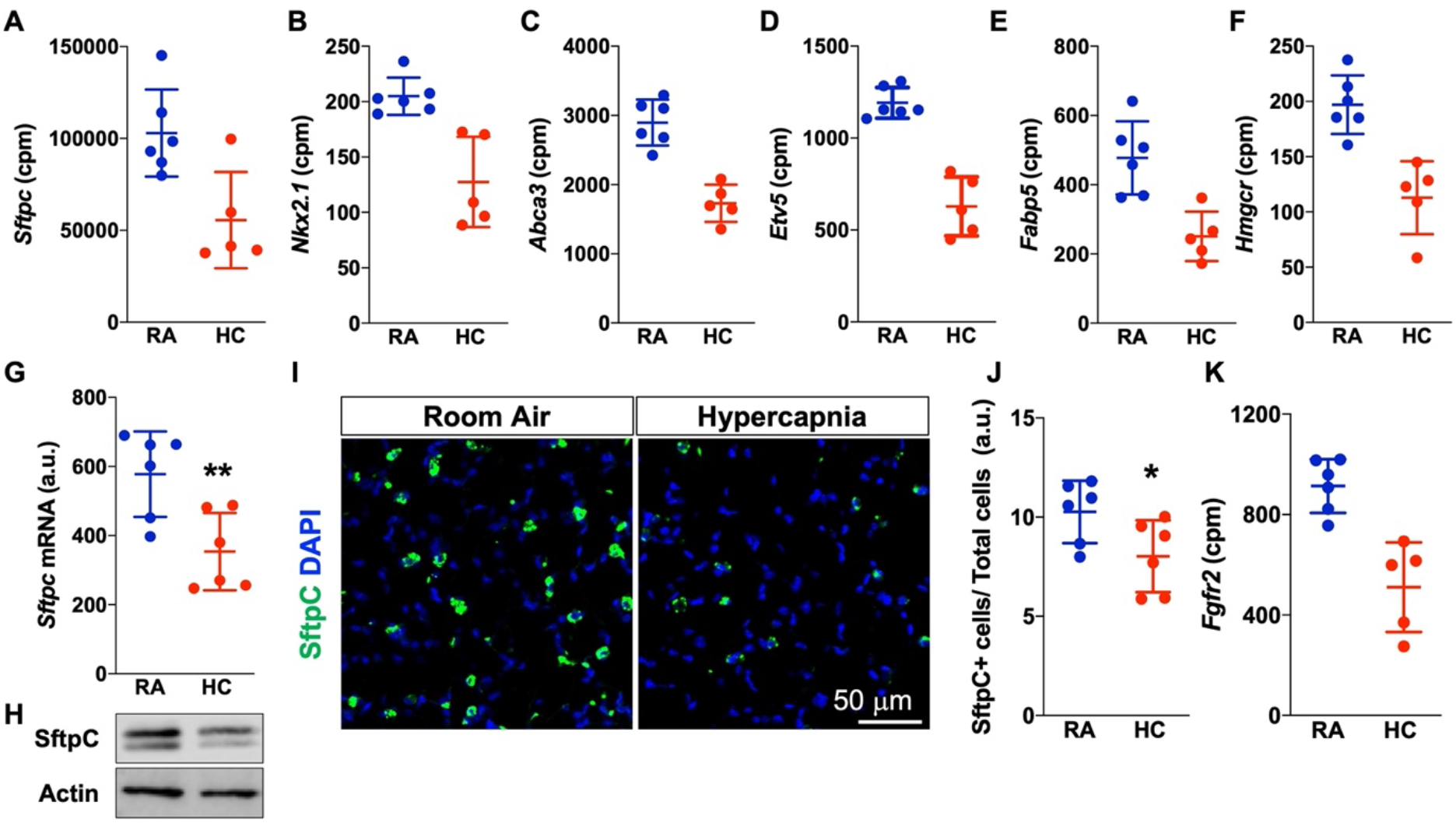
Transcriptomic analysis of AT2 cells shows that hypercapnia decreases the expression of AT2 markers. (**A-F**) Bulk RNASeq was performed on flow cytometry sorted AT2 cells from mice breathing room air (RA, n=6) or exposed to 10% CO_2_ (HC, n=5). Hypercapnia decreases the expression of AT2 markers. Regulation by hypercapnia of the expression of selected DEG markers (FDR *q*< 0.05) of AT2 cell regulated. (**G and H**) Quantification of SftpC expression in AT2 cells isolated from mice exposed to 10% CO_2_. (G) mRNA. n=6 mice. (H) A representative Western blot is shown. Actin was used as a loading control. (**I and J**) Quantification of SftpC^+^-AT2 cells in the alveolar region of adult mouse lung from mice breathing RA or exposed to HC for 21 days. n=6. (**K**) Hypercapnia decreases the expression of *Fgfr2* in sorted AT2 cells as analyzed by RNASeq. (FDR *q*< 0.05). \**P*<0.05; \*\**P*<0.01. Student’s *t*-test.

**Fig. S3.**
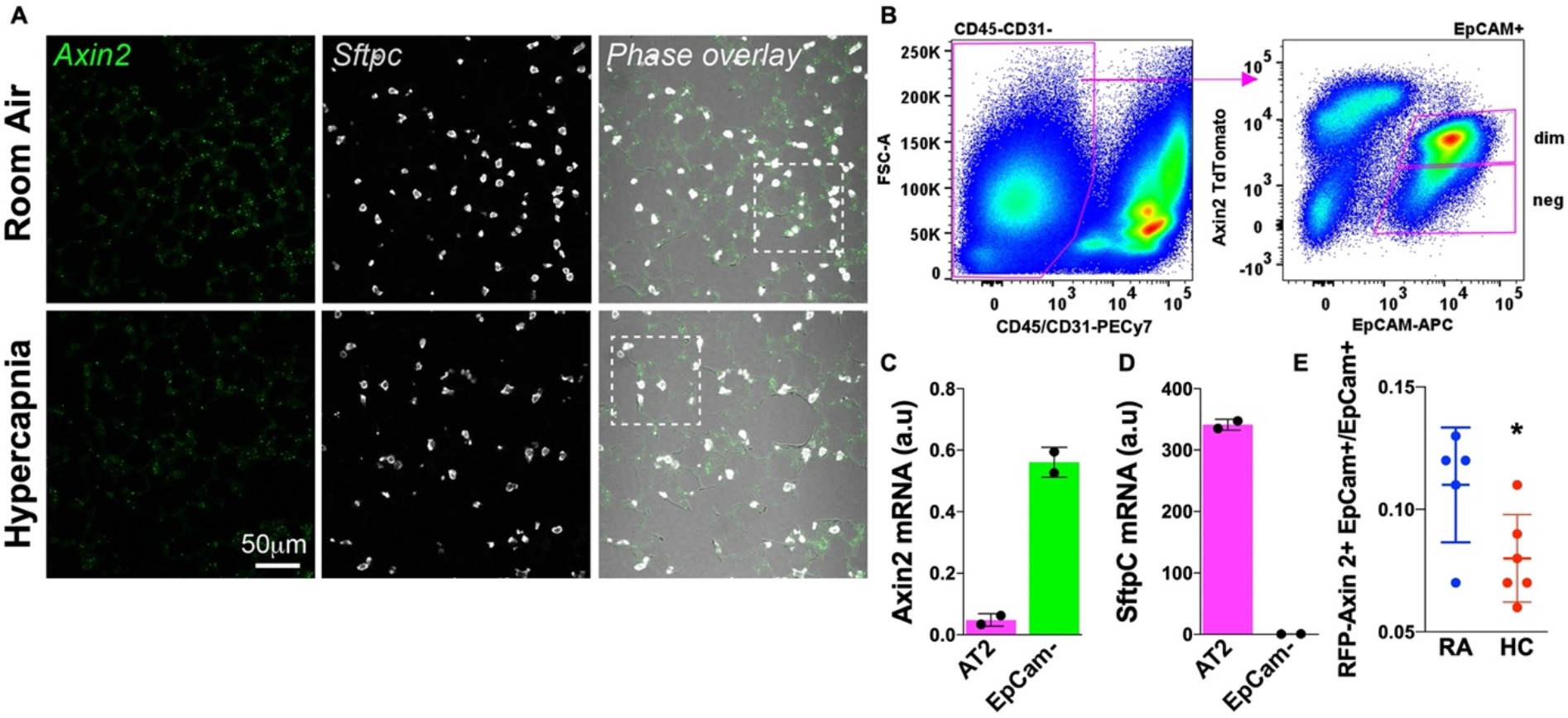
Hypercapnia decreases the number of Wnt-responsive AT2 cells. (**A**) Low magnification of *in situ* RNA hybridization showing the decreased number of Axin2^+^-AT2 cells in mice exposed to hypercapnia showed in Fig. 3B. (**B**) Gating strategy used to isolate Axin2^+^-AT2 cells (dim) from *Axin2^CreERT2-TdTom^* mice. (**C-D**) Expression of *Axin2* and *Sftpc* mRNA by RT-qPCR in AT2 and EpCam^-^ cells isolated from *Axin2^CreERT2-TdTom^* mice.

**Fig S4.**
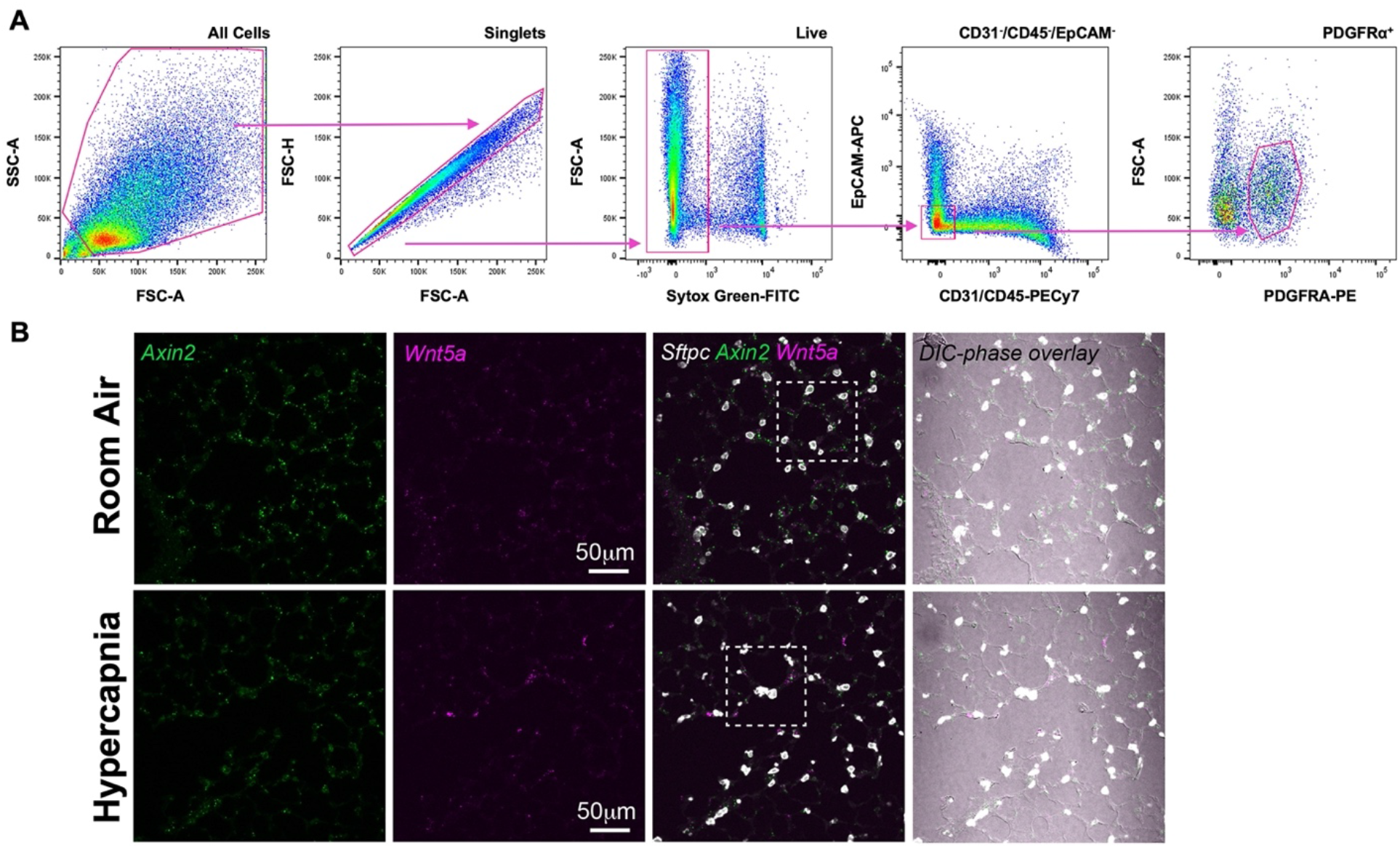
Hypercapnia alters Wnt expression in PDGFRα^+^ fibroblasts. (**A**) Gating strategy to isolate PDGFRα^+^ from mice. (**B**) Low magnification of *in situ* RNA hybridization showing the decreased number of *Axin2*^+^-AT2 cells in mice exposed to hypercapnia. Selected areas are showed in Fig. 4E.

**Fig S5.**
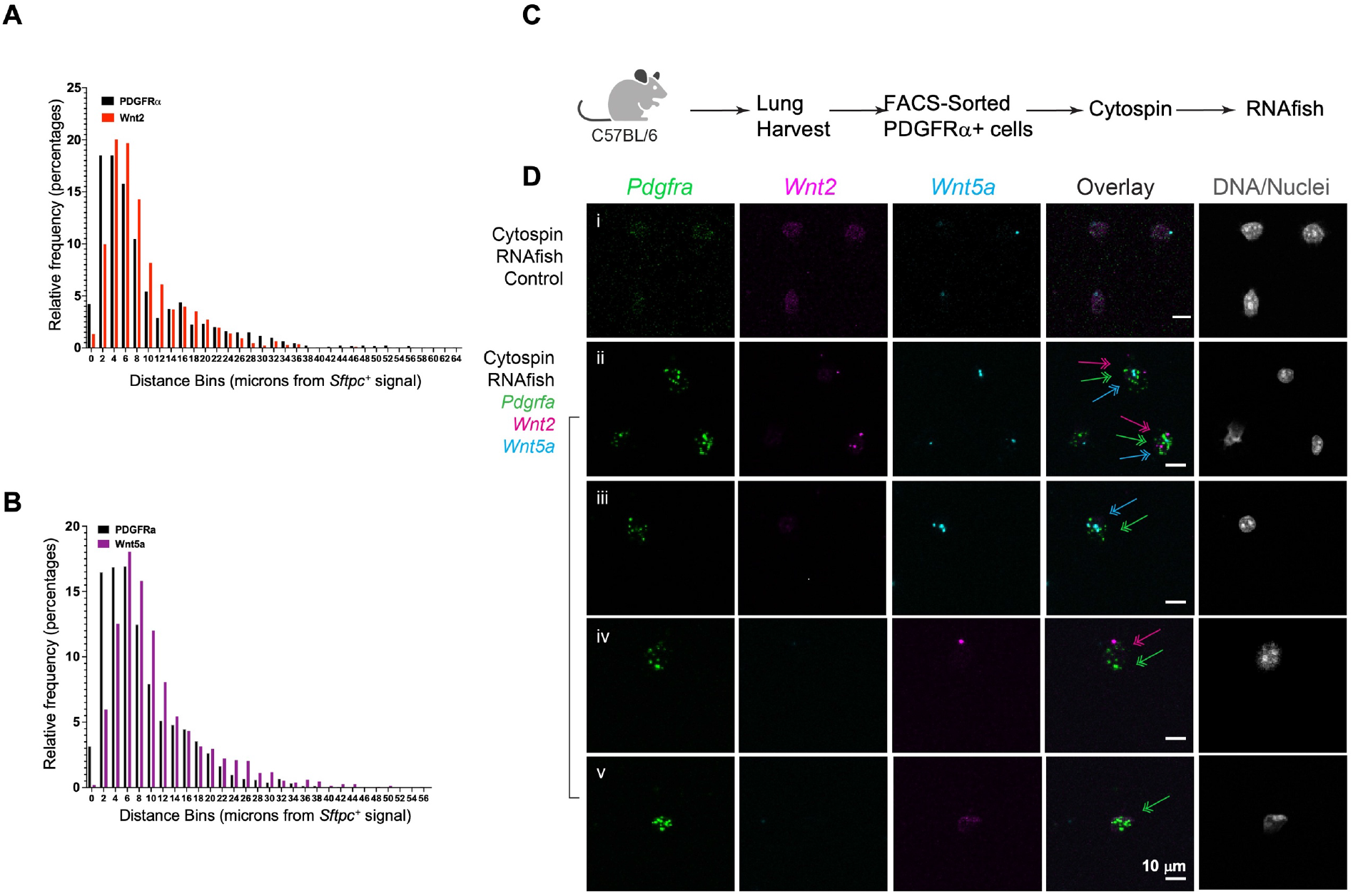
Single molecule RNAFISH analysis of PDGFRα^+^ lung fibroblasts. (**A and B**) Histograms for data showed in Fig. 6 A and B respectively. (**C**) Experimental design schematic for experiment in Fig. 6E. (**D**) Representative cytospin images for data in Fig 6E. PDGFRα-flow sorted fibroblasts subjected to RNAFISH with probes to *Pdgfra, Wnt2* and *Wnt5a* (ii-v, show different fields of view). Negative control also shown (i). Colored arrows correspond to respective probes and reveal *Pdgfra*+ cells single, double -positive or null for probes shown.

## REFERENCES

1. L. A. Dada, J. I. Sznajder, Mechanisms of pulmonary edema clearance during acute hypoxemic respiratory failure: role of the Na,K-ATPase. Crit Care Med 31, S248–252 (2003).

2. J. I. Sznajder, Alveolar Edema Must Be Cleared for the Acute Respiratory Distress Syndrome Patient to Survive. Am. J. Respir. Crit. Care Med. 163, 1293–1294 (2001).

3. A. Nalbandian, K. Sehgal, A. Gupta, M. V. Madhavan, C. McGroder, J. S. Stevens, J. R. Cook, A. S. Nordvig, D. Shalev, T. S. Sehrawat, N. Ahluwalia, B. Bikdeli, D. Dietz, C. Der-Nigoghossian, N. Liyanage-Don, G. F. Rosner, E. J. Bernstein, S. Mohan, A. A. Beckley, D. S. Seres, T. K. Choueiri, N. Uriel, J. C. Ausiello, D. Accili, D. E. Freedberg, M. Baldwin, A. Schwartz, D. Brodie, C. K. Garcia, M. S. V. Elkind, J. M. Connors, J. P. Bilezikian, D. W. Landry, E. Y. Wan, Post-acute COVID-19 syndrome. Nat Med 27, 601–615 (2021).

4. L. Morales-Quinteros, M. Camprubi-Rimblas, J. Bringue, L. D. Bos, M. J. Schultz, A. Artigas, The role of hypercapnia in acute respiratory failure. Intensive Care Med Exp 7, 39 (2019).

5. T. Barnes, V. Zochios, K. Parhar, Re-examining Permissive Hypercapnia in ARDS: A Narrative Review. Chest 154, 185–195 (2018).

6. M. Shigemura, E. Lecuona, J. I. Sznajder, Effects of hypercapnia on the lung. J Physiol, (2017).

7. I. Vadasz, R. D. Hubmayr, N. Nin, P. H. Sporn, J. I. Sznajder, Hypercapnia: a nonpermissive environment for the lung. Am J Respir Cell Mol Biol 46, 417–421 (2012).

8. C. U. Vohwinkel, E. Lecuona, H. Sun, N. Sommer, I. Vadasz, N. S. Chandel, J. I. Sznajder, Elevated CO(2) levels cause mitochondrial dysfunction and impair cell proliferation. J Biol Chem 286, 37067–37076 (2011).

9. A. Bharat, N. Graf, A. Mullen, J. Kanter, A. C. Andrei, P. H. Sporn, M. M. DeCamp, J. I. Sznajder, Pleural Hypercarbia After Lung Surgery Is Associated With Persistent Alveolopleural Fistulae. Chest 149, 220–227 (2016).

10. A. Bharat, M. Angulo, H. Sun, M. Akbarpour, A. Alberro, Y. Cheng, M. Shigemura, S. Berdnikovs, L. C. Welch, J. A. Kanter, High CO2 levels impair lung wound healing. Am J Respir Cell Mol Biol 63, 244–254 (2020).

11. T. J. Desai, D. G. Brownfield, M. A. Krasnow, Alveolar progenitor and stem cells in lung development, renewal and cancer. Nature 507, 190–194 (2014).

12. M. Aspal, R. L. Zemans, Mechanisms of ATII-to-ATI Cell Differentiation during Lung Regeneration. Int J Mol Sci 21, (2020).

13. C. E. Barkauskas, M. J. Cronce, C. R. Rackley, E. J. Bowie, D. R. Keene, B. R. Stripp, S. H. Randell, P. W. Noble, B. L. Hogan, Type 2 alveolar cells are stem cells in adult lung. J Clin invest 123, 3025–3036 (2013).

14. B. L. Hogan, C. E. Barkauskas, H. A. Chapman, J. A. Epstein, R. Jain, C. C. Hsia, L. Niklason, E. Calle, A. Le, S. H. Randell, J. Rock, M. Snitow, M. Krummel, B. R. Stripp, T. Vu, E. S. White, J. A. Whitsett, E. E. Morrisey, Repair and regeneration of the respiratory system: complexity, plasticity, and mechanisms of lung stem cell function. Cell Stem Cell 15, 123–138 (2014).

15. M. C. Basil, E. E. Morrisey, Lung regeneration: a tale of mice and men. Semin Cell Dev Biol, (2019).

16. W. J. Zacharias, D. B. Frank, J. A. Zepp, M. P. Morley, F. A. Alkhaleel, J. Kong, S. Zhou, E. Cantu, E. E. Morrisey, Regeneration of the lung alveolus by an evolutionarily conserved epithelial progenitor. Nature 555, 251–255 (2018).

17. A. S. Flozak, A. P. Lam, S. Russell, M. Jain, O. N. Peled, K. A. Sheppard, R. Beri, G. M. Mutlu, G. R. Budinger, C. J. Gottardi, Beta-catenin/T-cell factor signaling is activated during lung injury and promotes the survival and migration of alveolar epithelial cells. J Biol Chem 285, 3157–3167 (2010).

18. D. B. Frank, T. Peng, J. A. Zepp, M. Snitow, T. L. Vincent, I. J. Penkala, Z. Cui, M. J. Herriges, M. P. Morley, S. Zhou, M. M. Lu, E. E. Morrisey, Emergence of a Wave of Wnt Signaling that Regulates Lung Alveologenesis by Controlling Epithelial Self-Renewal and Differentiation. Cell Rep 17, 2312–2325 (2016).

19. A. N. Nabhan, D. G. Brownfield, P. B. Harbury, M. A. Krasnow, T. J. Desai, Singlecell Wnt signaling niches maintain stemness of alveolar type 2 cells. Science 359, 1118–1123 (2018).

20. J. H. Lee, T. Tammela, M. Hofree, J. Choi, N. D. Marjanovic, S. Han, D. Canner, K. Wu, M. Paschini, D. H. Bhang, T. Jacks, A. Regev, C. F. Kim, Anatomically and Functionally Distinct Lung Mesenchymal Populations Marked by Lgr5 and Lgr6. Cell 170, 1149–1163.e1112 (2017).

21. J. A. Zepp, W. J. Zacharias, D. B. Frank, C. A. Cavanaugh, S. Zhou, M. P. Morley, E. E. Morrisey, Distinct Mesenchymal Lineages and Niches Promote Epithelial Self-Renewal and Myofibrogenesis in the Lung. Cell 170, 1134–1148 e1110 (2017).

22. N. D. Magnani, L. A. Dada, M. A. Queisser, P. L. Brazee, L. C. Welch, K. R. Anekalla, G. Zhou, O. Vagin, A. V. Misharin, G. R. S. Budinger, K. Iwai, A. J. Ciechanover, J. I. Sznajder, HIF and HOIL-1L-mediated PKCzeta degradation stabilizes plasma membrane Na,K-ATPase to protect against hypoxia-induced lung injury. Proc Natl Acad Sci U S A 114, E10178–E10186 (2017).

23. P. L. Brazee, L. Morales-Nebreda, N. D. Magnani, J. G. Garcia, A. V. Misharin, K. M. Ridge, G. S. Budinger, K. Iwai, L. A. Dada, J. I. Sznajder, Linear ubiquitin assembly complex regulates lung epithelial–driven responses during influenza infection. J Clin Invest 130, 1301–1314 (2020).

24. A. Briva, I. Vadasz, E. Lecuona, L. C. Welch, J. Chen, L. A. Dada, H. E. Trejo, V. Dumasius, Z. S. Azzam, P. M. Myrianthefs, D. Batlle, Y. Gruenbaum, J. I. Sznajder, High CO(2) Levels Impair Alveolar Epithelial Function Independently of pH. PLoS ONE 2, e1238 (2007).

25. I. Vadasz, L. A. Dada, A. Briva, H. E. Trejo, L. C. Welch, J. Chen, P. T. Toth, E. Lecuona, L. A. Witters, P. T. Schumacker, N. S. Chandel, W. Seeger, J. I. Sznajder, AMP-activated protein kinase regulates CO2-induced alveolar epithelial dysfunction in rats and human cells by promoting Na,K-ATPase endocytosis. J Clin Invest 118, 752–762 (2008).

26. A. V. Misharin, C. M. Cuda, R. Saber, J. D. Turner, A. K. Gierut, G. K. Haines, 3rd, S. Berdnikovs, A. Filer, A. R. Clark, C. D. Buckley, G. M. Mutlu, G. R. Budinger, H. Perlman, Nonclassical Ly6C(-) monocytes drive the development of inflammatory arthritis in mice. Cell Rep 9, 591–604 (2014).

27. P. A. Reyfman, J. M. Walter, N. Joshi, K. R. Anekalla, A. C. McQuattie-Pimentel, S. Chiu, R. Fernandez, M. Akbarpour, C. I. Chen, Z. Ren, R. Verma, H. Abdala-Valencia, K. Nam, M. Chi, S. Han, F. J. Gonzalez-Gonzalez, S. Soberanes, S. Watanabe, K. J. N. Williams, A. S. Flozak, T. T. Nicholson, V. K. Morgan, D. R. Winter, M. Hinchcliff, C. L. Hrusch, R. D. Guzy, C. A. Bonham, A. I. Sperling, R. Bag, R. B. Hamanaka, G. M. Mutlu, A. V. Yeldandi, S. A. Marshall, A. Shilatifard, L. A. N. Amaral, H. Perlman, J. I. Sznajder, A. C. Argento, C. T. Gillespie, J. Dematte, M. Jain, B. D. Singer, K. M. Ridge, A. P. Lam, A. Bharat, S. M. Bhorade, C. J. Gottardi, G. R. S. Budinger, A. V. Misharin, Single-Cell Transcriptomic Analysis of Human Lung Provides Insights into the Pathobiology of Pulmonary Fibrosis. Am J Respir Crit Care Med, (2018).

28. J. Choi, J. E. Park, G. Tsagkogeorga, M. Yanagita, B. K. Koo, N. Han, J. H. Lee, Inflammatory Signals Induce AT2 Cell-Derived Damage-Associated Transient Progenitors that Mediate Alveolar Regeneration. Cell Stem Cell 27, 366–382.e367 (2020).

29. Z. Zhang, K. Newton, S. K. Kummerfeld, J. Webster, D. S. Kirkpatrick, L. Phu, J. Eastham-Anderson, J. Liu, W. P. Lee, J. Wu, Transcription factor Etv5 is essential for the maintenance of alveolar type II cells. Proc Natl Acad Sci U S A 114, 3903–3908 (2017).

30. B. Treutlein, D. G. Brownfield, A. R. Wu, N. F. Neff, G. L. Mantalas, F. H. Espinoza, T. J. Desai, M. A. Krasnow, S. R. Quake, Reconstructing lineage hierarchies of the distal lung epithelium using single-cell RNA-seq. Nature 509, 371–375 (2014).

31. S. J. Dorry, B. O. Ansbro, D. M. Ornitz, G. M. Mutlu, R. D. Guzy, FGFR2 Is Required for AEC2 Homeostasis and Survival after Bleomycin-induced Lung Injury. Am J Respir Cell Mol Biol 62, 608–621 (2020).

32. G. I. Balasooriya, M. Goschorska, E. Piddini, E. L. Rawlins, FGFR2 is required for airway basal cell self-renewal and terminal differentiation. Development 144, 1600–1606 (2017).

33. A. A. Raslan, J. K. Yoon, WNT signaling in lung repair and regeneration. Molecules and cells 43, 774 (2020).

34. J. Larraguibel, A. R. Weiss, D. J. Pasula, R. S. Dhaliwal, R. Kondra, T. J. Van Raay, Wnt ligand-dependent activation of the negative feedback regulator Nkd1. Mol Biol Cell 26, 2375–2384 (2015).

35. C. Bänziger, D. Soldini, C. Schütt, P. Zipperlen, G. Hausmann, K. Basler, Wntless, a conserved membrane protein dedicated to the secretion of Wnt proteins from signaling cells. Cell 125, 509–522 (2006).

36. C. M. Li, C. E. Kim, A. A. Margolin, M. Guo, J. Zhu, J. M. Mason, T. W. Hensle, V. V. Murty, P. E. Grundy, E. R. Fearon, V. D’Agati, J. D. Licht, B. Tycko, CTNNB1 mutations and overexpression of Wnt/beta-catenin target genes in WT1-mutant Wilms’ tumors. Am J Pathol 165, 1943–1953 (2004).

37. E.-h. Jho, T. Zhang, C. Domon, C.-K. Joo, J.-N. Freund, F. Costantini, Wnt/β-catenin/Tcf signaling induces the transcription of Axin2, a negative regulator of the signaling pathway. Mol Cell Biol 22, 1172–1183 (2002).

38. M. Shigemura, E. Lecuona, M. Angulo, L. A. Dada, M. B. Edwards, L. C. Welch, S. M. Casalino-Matsuda, P. H. S. Sporn, I. Vadasz, I. T. Helenius, G. A. Nader, Y. Gruenbaum, K. Sharabi, E. Cummins, C. Taylor, A. Bharat, C. J. Gottardi, G. J. Beitel, N. Kaminski, G. R. S. Budinger, S. Berdnikovs, J. I. Sznajder, Elevated CO2 regulates the Wnt signaling pathway in mammals, Drosophila melanogaster and Caenorhabditis elegans. Sci Rep 9, 18251 (2019).

39. H. Baarsma, M. Königshoff, ‘WNT-er is coming’: WNT signalling in chronic lung diseases. Thorax 72, 746–759 (2017).

40. X. Wu, E. M. van Dijk, J. P. Ng-Blichfeldt, I. S. T. Bos, C. Ciminieri, M. Königshoff, L. E. M. Kistemaker, R. Gosens, Mesenchymal WNT-5A/5B Signaling Represses Lung Alveolar Epithelial Progenitors. Cells 8, (2019).

41. R. Najdi, K. Proffitt, S. Sprowl, S. Kaur, J. Yu, T. M. Covey, D. M. Virshup, M. L. Waterman, A uniform human Wnt expression library reveals a shared secretory pathway and unique signaling activities. Differentiation 84, 203–213 (2012).

42. M. F. Miller, E. D. Cohen, J. E. Baggs, M. M. Lu, J. B. Hogenesch, E. E. Morrisey, Wnt ligands signal in a cooperative manner to promote foregut organogenesis. Proc Natl Acad Sci U S A 109, 15348–15353 (2012).

43. M. E. Rieger, B. Zhou, N. Solomon, M. Sunohara, C. Li, C. Nguyen, Y. Liu, J. H. Pan, P. Minoo, E. D. Crandall, S. L. Brody, M. Kahn, Z. Borok, p300/beta-Catenin Interactions Regulate Adult Progenitor Cell Differentiation Downstream of WNT5a/Protein Kinase C (PKC). J Biol Chem 291, 6569–6582 (2016).

44. A. S. Flozak, A. P. Lam, C. J. Gottardi, A Simple Method to Assess Abundance of the β-Catenin Signaling Pool in Cells. Methods Mol Biol 1481, 49–60 (2016).

45. N. Kramer, J. Schmöllerl, C. Unger, H. Nivarthi, A. Rudisch, D. Unterleuthner, M. Scherzer, A. Riedl, M. Artaker, I. Crncec, D. Lenhardt, T. Schwarz, B. Prieler, X. Han, M. Hengstschläger, J. Schüler, R. Eferl, R. Moriggl, W. Sommergruber, H. Dolznig, Autocrine WNT2 signaling in fibroblasts promotes colorectal cancer progression. Oncogene 36, 5460–5472 (2017).

46. Y. S. Jung, S. Jun, S. H. Lee, A. Sharma, J. I. Park, Wnt2 complements Wnt/β-catenin signaling in colorectal cancer. Oncotarget 6, 37257–37268 (2015).

47. A. J. Harwood, Regulation of GSK-3: a cellular multiprocessor. Cell 105, 821–824 (2001).

48. N. Harada, Y. Tamai, T. Ishikawa, B. Sauer, K. Takaku, M. Oshima, M. M. Taketo, Intestinal polyposis in mice with a dominant stable mutation of the beta-catenin gene. EMBO J 18, 5931–5942 (1999).

49. H. A. Baarsma, W. Skronska-Wasek, K. Mutze, F. Ciolek, D. E. Wagner, G. John-Schuster, K. Heinzelmann, A. Günther, K. R. Bracke, M. Dagouassat, Noncanonical WNT-5A signaling impairs endogenous lung repair in COPD. J Exp Med 214, 143–163 (2017).

50. Z. Steinhart, S. Angers, Wnt signaling in development and tissue homeostasis. Development 145, dev146589 (2018).

51. H. A. Chapman, X. Li, J. P. Alexander, A. Brumwell, W. Lorizio, K. Tan, A. Sonnenberg, Y. Wei, T. H. Vu, Integrin alpha6beta4 identifies an adult distal lung epithelial population with regenerative potential in mice. J Clin Invest 121, 2855–2862 (2011).

52. M. Shigemura, E. Lecuona, M. Angulo, T. Homma, D. A. Rodriguez, F. J. Gonzalez-Gonzalez, L. C. Welch, L. Amarelle, S. J. Kim, N. Kaminski, G. R. S. Budinger, J. Solway, J. I. Sznajder, Hypercapnia increases airway smooth muscle contractility via caspase-7-mediated miR-133a-RhoA signaling. Sci Transl Med 10, (2018).

53. K. L. Gates, H. A. Howell, A. Nair, C. U. Vohwinkel, L. C. Welch, G. J. Beitel, A. R. Hauser, J. I. Sznajder, P. H. Sporn, Hypercapnia impairs lung neutrophil function and increases mortality in murine pseudomonas pneumonia. Am J Respir Cell Mol Biol 49, 821–828 (2013).

54. A. Jaitovich, M. Angulo, E. Lecuona, L. A. Dada, L. C. Welch, Y. Cheng, G. Gusarova, E. Ceco, C. Liu, M. Shigemura, High CO2 levels cause skeletal muscle atrophy via AMP-activated kinase (AMPK), FoxO3a protein, and muscle-specific Ring finger protein 1 (MuRF1). J Biol Chem 290, 9183–9194 (2015).

55. K. Hasegawa, A. Sato, K. Tanimura, K. Uemasu, Y. Hamakawa, Y. Fuseya, S. Sato, S. Muro, T. Hirai, Fraction of MHCII and EpCAM expression characterizes distal lung epithelial cells for alveolar type 2 cell isolation. Respir Res 18, 150 (2017).

56. M. Lawrence, W. Huber, H. Pagès, P. Aboyoun, M. Carlson, R. Gentleman, M. T. Morgan, V. J. Carey, Software for computing and annotating genomic ranges. PLoS Comput Biol 9, e1003118 (2013).

57. M. D. Robinson, D. J. McCarthy, G. K. Smyth, edgeR: a Bioconductor package for differential expression analysis of digital gene expression data. Bioinformatics 26, 139–140 (2010).

58. D. J. McCarthy, Y. Chen, G. K. Smyth, Differential expression analysis of multifactor RNA-Seq experiments with respect to biological variation. Nucleic Acids Res 40, 4288–4297 (2012).

59. E. Eden, R. Navon, I. Steinfeld, D. Lipson, Z. Yakhini, GOrilla: a tool for discovery and visualization of enriched GO terms in ranked gene lists. BMC Bioinformatics 10, 48 (2009).

60. L. A. Dada, H. E. Trejo Bittar, L. C. Welch, O. Vagin, N. Deiss-Yehiely, A. M. Kelly, M. R. Baker, J. Capri, W. Cohn, J. P. Whitelegge, I. Vadasz, Y. Gruenbaum, J. I. Sznajder, High CO2 Leads to Na,K-ATPase Endocytosis via c-Jun Amino-Terminal Kinase-Induced LMO7b Phosphorylation. Mol Cell Biol 35, 3962–3973 (2015).

